# *De novo* designed transmembrane domains tune engineered receptor functions

**DOI:** 10.1101/2020.07.26.221598

**Authors:** Assaf Elazar, Nicholas J. Chandler, Ashleigh S. Davey, Jonathan Y. Weinstein, Julie V. Nguyen, Raphael Trenker, Ryan S. Cross, Misty R. Jenkins, Melissa J. Call, Matthew E. Call, Sarel J. Fleishman

**Author notes:** Equal senior and corresponding authors **Materials and Correspondence** Correspondence and requests for materials related to proMP design methods should be directed to S.J.F. Correspondence and requests for materials related to proMP structural analysis should be directed to M.J.C. Correspondence and requests for materials related to proCAR applications should be directed to M.E.C. Equal first authors (Assaf Elazar, Nicholas J. Chandler, Ashleigh S. Davey).

## Abstract

*De novo* designed receptor transmembrane domains (TMDs) present opportunities for precise control of cellular receptor functions. We developed a *de novo* design strategy for generating programmed membrane proteins (proMPs): single-pass α-helical TMDs that self-assemble through computationally defined and crystallographically validated interfaces. We used these proMPs to program specific oligomeric interactions into a chimeric antigen receptor (CAR) and found that both *in vitro* CAR T cell cytokine release and *in vivo* antitumor activity scaled linearly with the oligomeric state encoded by the receptor TMD, from monomers up to tetramers. All programmed CARs (proCARs) stimulated substantially lower T cell cytokine release relative to the commonly used CD28 TMD, which we show elevated cytokine release through lateral recruitment of the endogenous T cell costimulatory receptor CD28. Precise design using orthogonal and modular TMDs thus provides a new way to program receptor structure and predictably tune activity for basic or applied synthetic biology.

## Introduction

Interactions among cell-surface receptors play central roles in determining complex structures and controlling signal propagation. In immune receptors (Berry and Call, 2017; Dong et al., 2019), death receptors (Fu et al., 2016; Pan et al., 2019) and growth factor receptors (Arkhipov et al., 2013; Endres et al., 2013; Fleishman et al., 2002), the transmembrane domains (TMDs) govern key interactions involved in assembly, activation and high-order clustering. Control over the specificity, stability, geometry and oligomeric state of these interactions is therefore highly desirable both for mechanistic studies of natural receptors and in the engineering of synthetic receptors. Accurate control, however, is difficult to achieve using natural TMDs that have likely been evolutionarily selected for a degree of flexibility in these very attributes (Matthews et al., 2006). The importance of high precision in receptor engineering has come into particularly sharp focus with the clinical adoption of cancer immunotherapies using targeted chimeric antigen receptors (CARs) (Eshhar et al., 1993; June et al., 2018; Majzner and Mackall, 2019; Salter et al., 2018) to endow T cells with potent antitumor activity. Controlling functional outputs from these engineered single-chain immune receptors poses significant challenges in balancing antitumor CAR T cell activity against toxicities associated with high inflammatory cytokine release, known as cytokine release syndrome (CRS) (Gutierrez et al., 2018; Maus et al., 2020; Morgan et al., 2010; Morris et al., 2021).

The modular domain organization of CARs offers a prime example of a synthetic cellular system in which customizable sequences exert control over receptor structure and function. Current-generation CARs comprise an antibody single-chain variable fragment (scFv) domain for tumor antigen binding, a spacer or hinge domain for length and flexibility, a TMD controlling membrane integration and expression levels, and intracellular costimulation and activation domains that provide signals for proliferation, survival and activation of T cell effector functions. Efforts to imbue CARs with optimal signaling properties have probed all of these domains in one way or another (Alabanza et al., 2017; Balakrishnan et al., 2019; Feucht et al., 2019; Hartl et al., 2020; James, 2018; Liu et al., 2015; Majzner et al., 2020; Mata and Gottschalk, 2019; Rafiq et al., 2020; Wu et al., 2020). The TMDs, however, have received little attention in systematic studies of CAR design. For convenience, most incorporate the TMD sequence of the protein from which the adjacent hinge or signalling domains were derived; that is, most commonly from endogenous T cell proteins such as CD4, CD8, CD28 or the T cell receptor (TCR)-associated CD3ζ chain. At least some of these TMD sequences can engage in molecular interactions that drive self-association and/or assembly with the essential T cell proteins from which they were derived and thereby impact CAR expression and functions in ways that reduce control over signalling outcomes (Bridgeman et al., 2010, 2014; Call et al., 2006; Cosson et al., 1991; Hennecke and Cosson, 1993; Leddon et al., 2020; Muller et al., 2021). Thus, the involvement of these natural immune-receptor TMDs in native T cell signaling hampers rational design of CARs with predictable properties.

We set out to define the relationships between TMD structure, CAR oligomeric state and signalling in CAR T cells by designing completely new TMDs with programmable self-association features and minimal risk of cross-talk with native T cell components. Despite significant recent progress (Barth and Senes, 2016; Korendovych and DeGrado, 2020), the limitations of membrane-protein (MP) atomistic calculations have restricted *de novo* a-helical MP design studies to highly predictable and rigid coiled-coil motifs (Joh et al., 2014; Lu et al., 2018) that, while stabilising them, limited their usefulness as receptor TMDs. By contrast, we recently described an *ab initio* Rosetta atomistic modelling strategy (Weinstein et al., 2019) that uses a new energy function with experimentally determined membrane-solvation terms for each amino acid. This modeling strategy accurately predicts the structure of single-spanning sequences known to self-assemble (Elazar et al., 2016a, 2016b). Here, we introduce a new strategy to *de novo* design proMPs (**pro**grammable **M**embrane **P**roteins) resulting in completely new sequences that form TM homo-oligomers of defined geometry and order that can be used to program cell-surface receptor structure. We used these proMPs to generate proCAR (**pro**grammed **CAR**) constructs and found that they endowed T cells with *in vivo* functional potency that scaled linearly with oligomeric state. ProCARs also maintained significantly lower inflammatory cytokine release compared to an otherwise identical CAR containing the natural CD28 TMD, a property that may have safety benefits in clinical applications (Alabanza et al., 2017; Brudno et al., 2020; Morris et al., 2021; Rafiq et al., 2020; Ying et al., 2019). Our results shed new light on the importance of precision in engineered receptor structure and intermolecular associations for optimal CAR T activity and provide new design tools that may be useful for developing cellular immunotherapies with optimal safety and efficacy profiles.

## Results

### Atomically precise *de novo* designed TMDs

In our initial design approach (**Figure 1a**), each design trajectory started from two fully symmetric and extended chains of 24 amino acids encoding either poly-Val or poly-Ala (Supplemental Movie). In a first, coarse-grained modelling step, backbone torsion angles were sampled from a database comprising 3 and 9 amino acid fragments from α-helical MPs, and the two chains were symmetrically docked against one another with an energy term that disfavoured large crossing angles (**Supplementary Eq. 1**) (Bowie, 1997; Weinstein et al., 2019). In a second, all-atom step, we refined the sequence and the structure through iterations of symmetric sequence optimisation, backbone minimisation, and rigid-body docking using the ref2015_memb atomistic energy function that is dominated by van der Waals packing, hydrogen bonding and amino acid lipophilicity (Weinstein et al., 2019). We noticed that the resulting sequences were overwhelmingly biased towards the large and flexible hydrophobic amino acid Leu (**Figure 1b**), as expected from the dominant role of lipophilicity in the ref2015_memb potential (Weinstein et al., 2019). Forward-folding *ab initio* structure-prediction calculations, however, indicated that the designs were prone to form multiple alternative low-energy dimers instead of the designed ones (**Figure S1a**). To mitigate the risk of misfolding due to the high Leu content, we introduced a sequence diversification step comprising 120 iterations of single-point mutation and energy relaxation while biasing the sequence composition to match that of natural TMDs (**Figure 1b**; **Eq. 2-3**). The resulting sequences were subjected to *ab initio* structure prediction calculations (Das et al., 2009) and this time, they converged to the design models (**Figure S1b**) and exhibited a large energy gap from undesired structures. Previous studies noted that natural TMDs are not optimised for thermodynamic stability (Faham et al., 2004). Our design simulations suggest that evolution might have selected sequence compositions to counter TMD misfolding.

**Figure 1:**
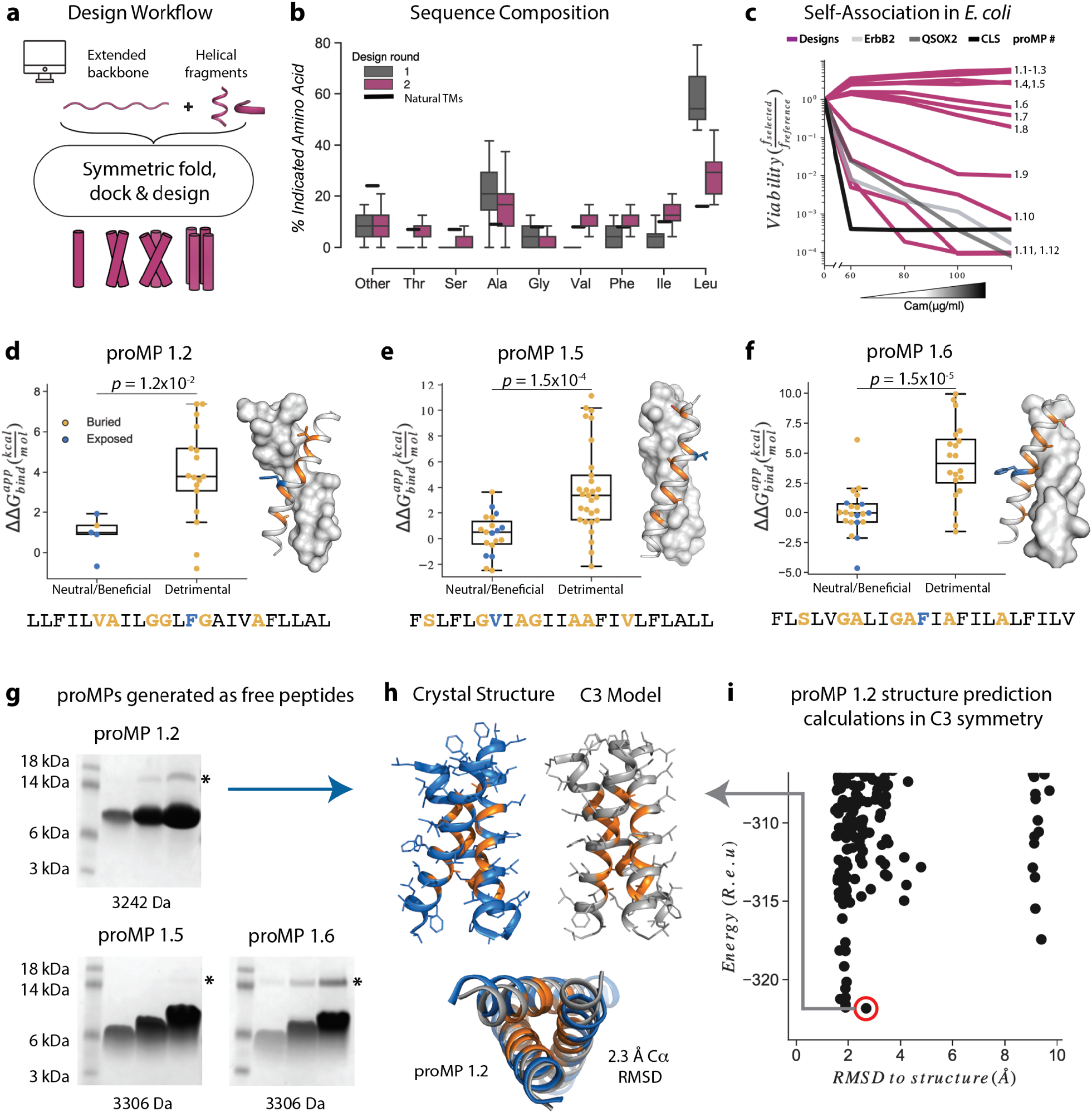
Learning the rules for programming self-associating MPs. (**a**) Rosetta fold, dock and design uses backbone fragments from natural MPs to construct symmetric, *de novo* architectures and a MP energy function(Weinstein et al., 2019) to optimise the amino acid sequence. (**b**) Round 1 designs were biased towards the hydrophobic amino acid Leu relative to naturally occurring TMDs. In round 2, we incorporated a sequence diversification step which conformed the amino acid propensities to those observed in natural TMDs. (**c**) The proMPs strongly self-associate in the *E. coli* inner membrane as evidenced by high viability in the dsTβL self-association assay(Elazar et al., 2016a). (**d-f**) Designed positions that are buried at the interface (orange) are more sensitive to mutation according to deep sequencing TβL analysis(Elazar et al., 2016a) (*y*-axis) than exposed positions (blue). Mutations are predicted to be detrimental or neutral/beneficial using computational mutation scanning of the model structures (Methods). Changes in self-association energies upon mutation are computed according to Eq. 9. (**g**) ProMPs produced as free peptides form SDS-stable homo-oligomers. SDS-PAGE samples containing approximately 15, 45 and 135 μg of peptide were heated to 95 °C for 1 minute and run under reducing conditions. (*) Indicates the position of a minor contaminant from the fusion protein used to generate proMP peptides (Methods). Molecular weight below each gel is for a monomer of the corresponding peptide sequence with additional N-terminal EPE and C-terminal RRLC flanking sequences (Methods). See additional examples in Figure S3. (**h, i**) The 2.55 Å resolution structure (blue ribbon) determined from crystals grown in monoolein lipid cubic phase (LCP) shows that proMP 1.2, designed to form a dimer, associates to form a trimer in a lipid bilayer environment. (**i**) Forward-folding *ab initio* prediction of proMP 1.2 in trimeric (C3) symmetry results in a model structure (**h**, gray ribbon) that is very close to the experimentally determined one.

Twelve designs were tested in the *E. coli* TOXCAT-β-lactamase (TβL) selection system (Elazar et al., 2016a; Langosch et al., 1996). In this dual-reporter system, survival on ampicillin and chloramphenicol reports on a design’s membrane-insertion and self-association propensity, respectively (**Figure S2**). Remarkably, most proMPs supported high survival (**Figure 1c**) and two-thirds survived even at the highest chloramphenicol concentration tested, indicating a self-association strength significantly greater than the TMD from the human receptor-tyrosine kinase HER2 (also known as ErbB2), which served as a positive control. Deep mutational scanning of mutant libraries showed that the sensitivity to mutations of most designs was consistent with interfacial versus exposed positions in the design models (**Figure 1d-f** and **Figure S3**), suggesting that they indeed assembled through the designed interfaces in the bacterial inner membrane.

Eight proMPs were produced recombinantly as free peptides and all exhibited electrophoretic mobility consistent with SDS- and heat-stable self-association (**Figure 1g** and **Figure S3**). The patterns of migration, however, were not uniform. Six proMPs had the apparent molecular weight of a dimer (*e*.*g*., proMP 1.5 and 1.6, **Figure 1g**) and exhibited reduced mobility as the peptide concentration was increased, similar to the SDS-stable (Lemmon et al., 1992) behaviour of the well-studied glycophorin A TMD (**Figure S3**). By contrast, the remaining two proMPs exhibited migration patterns that were independent of the sample concentration and had apparent molecular weights more consistent with oligomers larger than the designed dimers (proMP 1.2, **Figure 1g**; proMP 1.3, **Figure S3**). To establish the molecular structures of these designs, several were screened for crystallisation in monoolein lipid cubic phase, and the structure of proMP 1.2 was determined to 2.55 Å resolution (**Figure 1h** and **Figure S4**). While the positions involved in helix packing recapitulated the design model, this proMP indeed formed a trimer instead of the intended dimer. *Ab initio* structure prediction calculations in trimeric (C3) symmetry recapitulated the experimentally observed packing interface (RMSD 2.3 Å) (**Figure 1h-i**), demonstrating that it would have been possible to predict this outcome had we considered alternative oligomeric states during design calculations.

Based on this insight, we initiated a third design campaign to produce proMPs in a range of oligomeric states. We incorporated a final step in which *ab initio* structure prediction calculations (Weinstein et al., 2019) were performed in C2, C3 and C4 symmetries for every design. Only those proMPs that were predicted to form the target oligomeric state and none of the alternatives were selected for further analysis (**Figure 2a-d**). This strategy yielded two proMPs for which we obtained crystal structures confirming the target oligomeric state: a dimer with glycine-based packing interface similar to the motif observed in human glycophorin A (proMP C2.1; **Figure 2a-b, e-g** and **Figure S5**), and a trimer with an alanine-rich interface which, to the best of our knowledge, is novel in a membrane protein (proMP C3.1; **Figure 2c-d, h-k** and **Figure S6**). Interestingly, while two of the helices in the crystal structure of proMP C3.1 aligned well with the design model (**Figure 2i**), the third was in an antiparallel orientation (**Figure 2j**). Despite this arrangement, the six key interface alanine β-methyls were in near-identical positions to their counterparts in the fully parallel model (**Figure 2k**) leading us to suspect that the model is correct but the crystal lattice was enforcing the antiparallel binding mode of the third helix. To probe this possibility, we aligned the parallel model with the asymmetric unit seen in the crystal structure and generated crystallographic symmetry. The resulting model showed clashes for the third helix and indicated that the design model cannot be accommodated in the crystal lattice. The structure thus suggests that this proMP is unintentionally “reversible” in that one of the helices can form the intended packing mode in either orientation. While this feature is of interest from a design standpoint (Woodall et al., 2015), we note that only the fully parallel trimer depicted in the model can form in a biological system where the topology of a single-spanning TMD is constrained by the biosynthetic machinery in a type-I orientation.

**Figure 2:**
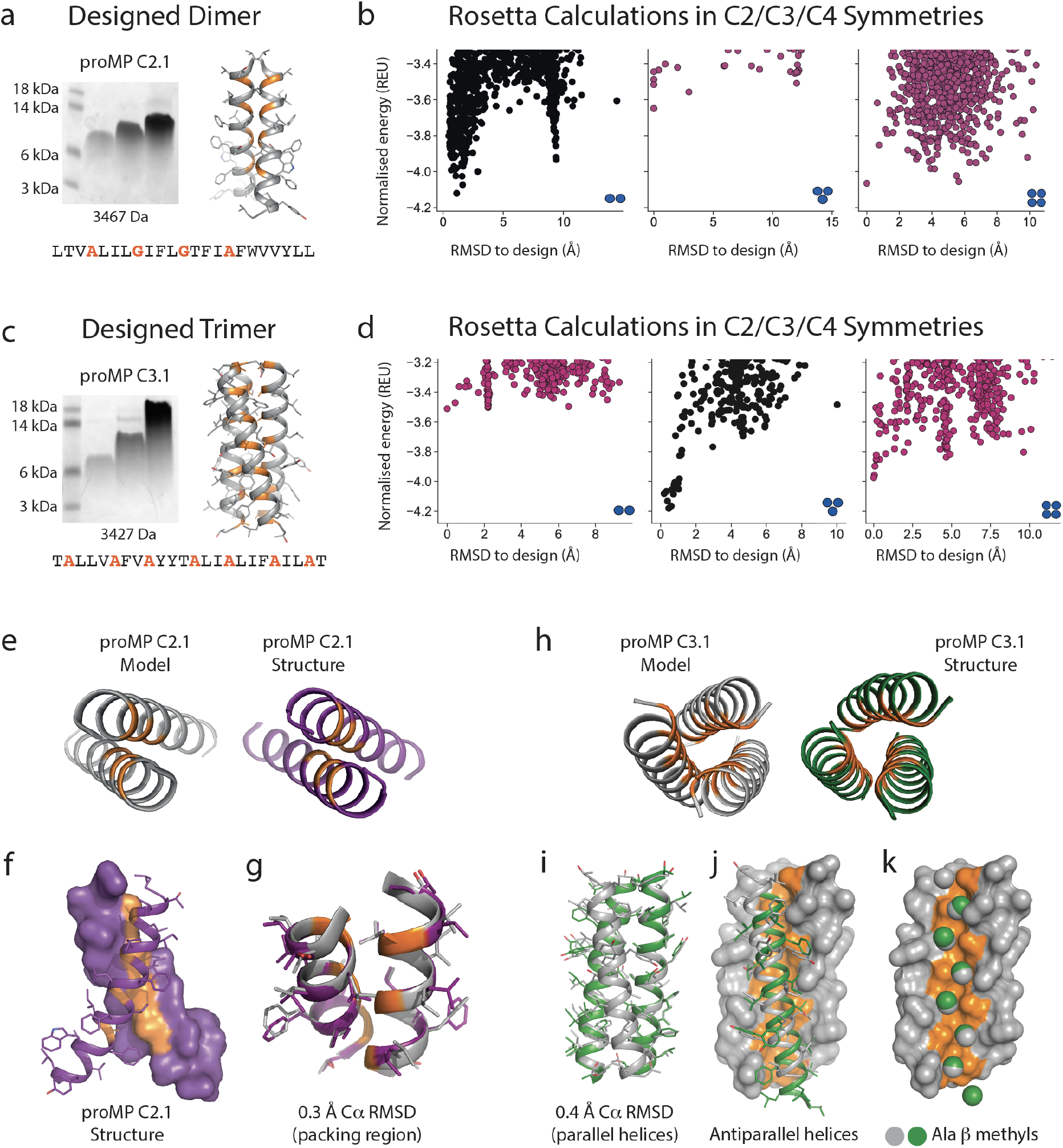
Designed MPs of defined structure and oligomeric state. (**a**) SDS-PAGE migration of proMP C2.1 is consistent with a dimer in gel-shift assays performed as in Figure 1. Design model and peptide sequence shown for reference. (**b**) Rosetta *ab initio* structure prediction calculations predict that proMP C2.1 preferentially forms a dimer. (**c**) ProMP C3.1 exhibits a novel Ala-dominated interface, and its migration pattern at high sample concentration suggests a complex larger than a dimer. Design model and peptide sequence shown for reference. (**d**) *Ab initio* calculations predict that it primarily forms a trimer. (**e-g**) The proMP C2.1 designed structure is atomically verified by a 2.7 Å crystal structure. Interfacial positions are marked in orange. (**h-k**) The crystallographic analysis of proMP C3.1 (3.5 Å resolution) reveals a trimer that is almost identical to the design, although one of the three helices in the trimer is antiparallel relative to the other two in the crystal lattice. Alignment of the structure and model (**i**) shows that the antiparallel helix (green) (**j**) positions Ala Cβ methyls that pack into the trimer through the designed interface (grey) (**k**).

We conclude that the sequence diversification and the computational selection of the oligomeric state described above provide a practical approach to implement negative-design principles that are critical for accurate *de novo* TMD design (Fleishman and Baker, 2012; Joh et al., 2014). These new insights will likely also be critical to design *de novo* hetero-oligomeric TMDs.

### Programmed CARs (proCARs) with defined oligomeric states

The availability of synthetic TMDs with defined structures provided an opportunity to address two key open questions in receptor engineering: What is the relationship between oligomeric state and functional output? And does the use of natural TMDs impart functional characteristics other than surface localization in the membrane? The hinge-TMD regions in all CARs used in FDA-approved CAR T cell products derive from CD8 or CD28 and drive disulphide-linked receptor homodimer formation (Fujiwara et al., 2020). However, the importance of the dimeric state for optimal CAR function is not well understood and alternative oligomeric forms such as trimers or tetramers have not been explored. Furthermore, both CD8 and CD28 TMDs have documented propensities to self-associate (Hennecke and Cosson, 1993; Leddon et al., 2020). Given the presence of both native receptors in CAR T cells, their use in CARs risks unintended interactions that could affect their expression and/or function. To program CARs that form specific oligomeric states and are insulated from confounding interactions with endogenous signalling proteins, we initially chose the crystallographically confirmed proMPs C2.1 and 1.2 to generate structurally programmed CARs (proCARs) that form dimers or trimers. These were termed proCAR-2 and proCAR-3, respectively (**Figure 3a**). We also designed a monomeric proMP that exhibited no chloramphenicol survival in dsTβL assays (**Figure S7**) and used it to produce a monomeric proCAR-1 in order to extend the structure-function study. Our proCAR designs incorporated an anti-HER2 scFv (FRP5 (Wels et al., 1992)) fused to the human CD8α hinge sequence, a proMP-derived TMD, the human CD28 costimulatory sequence and the human CD3ζ activating tail. Our reference CAR contained the human CD28 TMD for comparison, approximating a domain configuration that has been extensively studied *in vitro* and *in vivo* (Davenport et al., 2015, 2018; Haynes et al., 2002). In all proCAR constructs, a cysteine residue in the CD8α hinge that mediates disulphide-bonded dimer formation was mutated to alanine (**Figure 3a**) to ensure that the designed TMDs were the primary determinants of oligomeric state.

**Figure 3:**
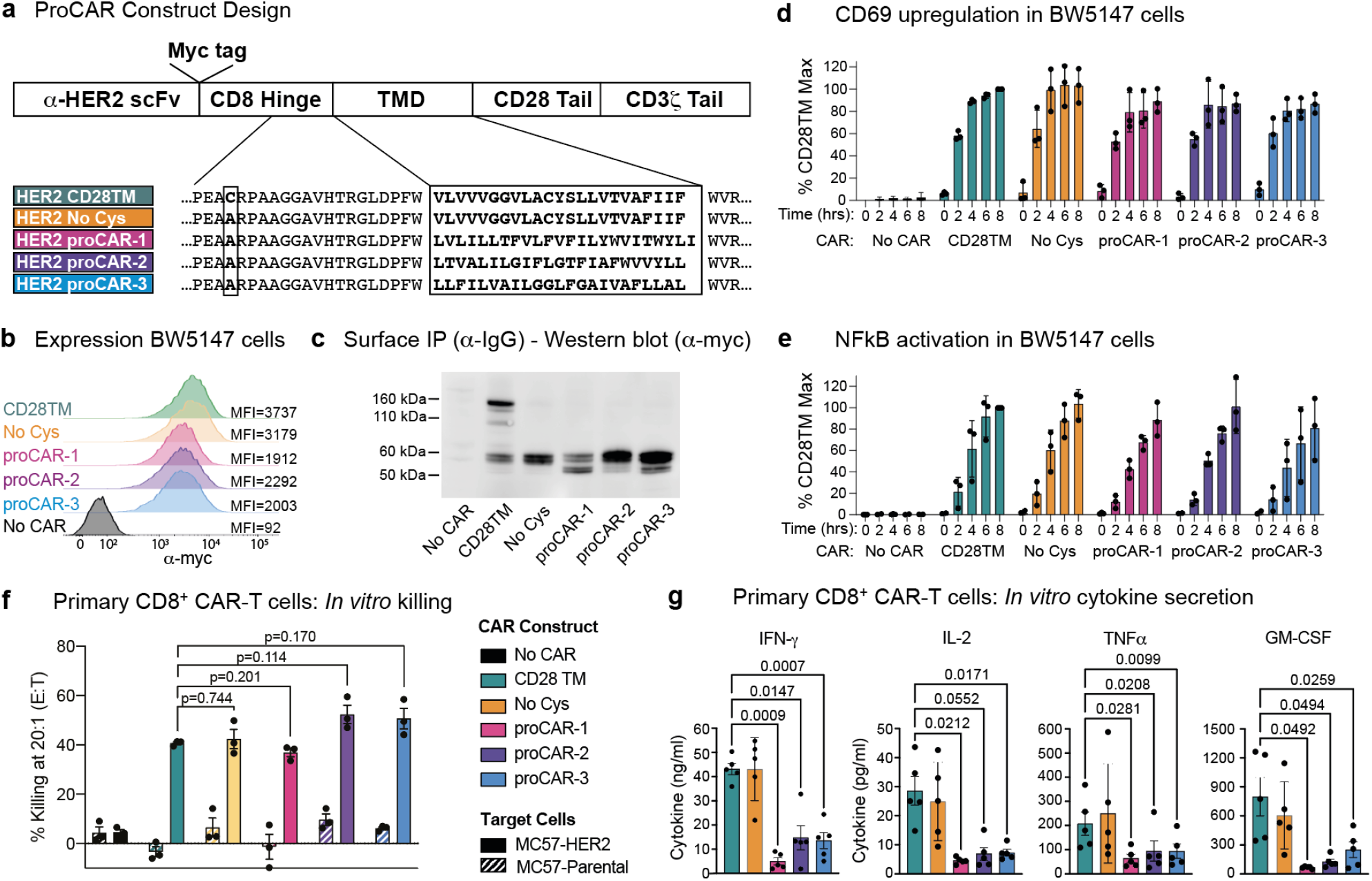
Construction and *in vitro* functional profiling of HER2-specific proCARs. **(a)** Schematic showing the domain organisation of the reference HER2-specific CAR constructs and modifications made to introduce proMP TMDs. Bold, boxed sequence indicates the human CD28 TMD in the reference CD28TM and no cys CARs and designed proMP sequences in the monomeric (proCAR-1), dimeric (proCAR-2) and trimeric (proCAR-3) receptors. **(b)** BW5147 murine thymoma cells stably expressing proCARs and a destabilised GFP NF-**κ**B reporter were surface labelled with anti-myc antibody and analysed by flow cytometry to assess surface expression levels. **(c)** Live cells from (b) were coated with polyclonal anti-IgG to bind CARs through the scFv domain and immunoprecipitated using protein G beads. Products were separated by non-reducing SDS-PAGE and immunoblotted using anti-myc antibody to visualise surface-expressed CAR proteins. Molecular weight of the unglycosylated CAR polypeptide is 55 kDa. **(d, e)** Cells from (b) were co-cultured with HER2+ SKBR3 human breast adenocarcinoma cells for the indicated times and analysed by flow cytometry for upregulation of activation marker CD69 **(d)** and GFP expression from the NF-**κ**B reporter **(e)**. All activation levels are normalised to the 8-hr time point in cells expressing the CD28TM CAR (% CD28TM Max). Bars represent the mean ± SD, and dots show the individual data points for three independent experiments. (**f**) Maximum target killing percentage at 20:1 effector to target ratio from 4hr ^51^Cr release assay. Bars show mean ± SEM with each data point representing an individual experiment (n=3). P-values determined from paired *t*-tests. (**g**) Cytokine production by primary mouse HER2 proCAR T cells following 24hr co-culture with MC57-HER2 target tumor cells. Bars show mean concentration ± SEM with each data point representing an individual experiment (n=5). Significance was determined from one-way ANOVA with multiple comparisons. Cytokine production on antigen-negative parental MC57 cells shown separately in Figure S9.

The HER2 proCARs and reference CD28TM constructs were retrovirally expressed in murine BW5147 thymoma cells. All constructs exhibited similar cell-surface levels (**Figure 3b**) and the reference CD28TM CAR formed disulphide-linked dimers while the cysteine mutant reference (No Cys) and proCARs did not (**Figure 3c)**. All CARs were competent to signal when co-cultured with HER2^+^ SKBR3 human breast adenocarcinoma cells (**Figure 3d-e**). When expressed in freshly isolated mouse CD8^+^ T cells (**Figure S8a-b)**, all CARs mediated antigen-dependent killing of MC57 mouse fibrosarcoma cells stably expressing HER2 *in vitro* (**Figure 3f**). Only small differences in killing potency were apparent, with proCAR-1 trending slightly less effective than the reference CARs and proCAR-2 and -3 trending slightly more effective. *In vitro* cytokine production (IFNγ, IL-2, TNFα and GM-CSF), on the other hand, was significantly lower in all proCARs, reduced by 2-10 fold on average (**Figure 3g**). This effect was not apparent in the CD28TM (No Cys) background and was therefore not due to loss of the CD8α hinge-region disulphide bond.

### The CD28 TMD enhances CAR-mediated cytokine release by associating with endogenous T cell CD28

The striking reduction in cytokine release in all of the proCARs led us to hypothesize that the higher levels of cytokine release in CD28 TMD-containing CARs depend primarily on CD28 sequence features rather than on CAR oligomeric state. The CD28 TMD contains a highly conserved polar YxxxxT motif that is similar to one that drives CD3ζ dimerization (Call et al., 2006) and is required for optimal dimerisation and surface expression of native CD28 (Leddon et al., 2020). A recent study showed that the CD28 YxxxxT sequence also causes CARs containing the CD28 TMD to physically associate with the CD28 protein in T cells (Muller et al., 2021), but the functional consequences of this association for CAR signaling have not been explored. We modeled this putative CD28 TM interface on the ζζ structure (Call et al., 2006) and noted that tyrosine, serine and threonine in the YSLLVT sequence could all contribute to an interhelical hydrogen-bonding network (**Figure 4a**). We therefore generated a CAR in which this sequence was mutated to **FA**LLV**V**, selectively eliminating the key hydrogen-bonding hydroxyl groups, for comparison to our proCAR and reference constructs. This CAR was well expressed as a disulphide-linked homodimer at the cell surface (**Figure 4b-c**) and generated primary mouse CD8^+^ CAR T cells whose ability to kill HER2^+^ target cells was unimpaired (**Figure 4d**). The CD28 TM mutant, however, induced lower levels of cytokine secretion (2-to 6-fold lower on average; **Figure 4d**) that were similar to those observed for the proCARs. We therefore concluded that the low-cytokine release seen in the proCAR T cells was likely due to the proCARs being insulated from interaction with endogenous T cell signalling proteins, primarily CD28.

**Figure 4.**
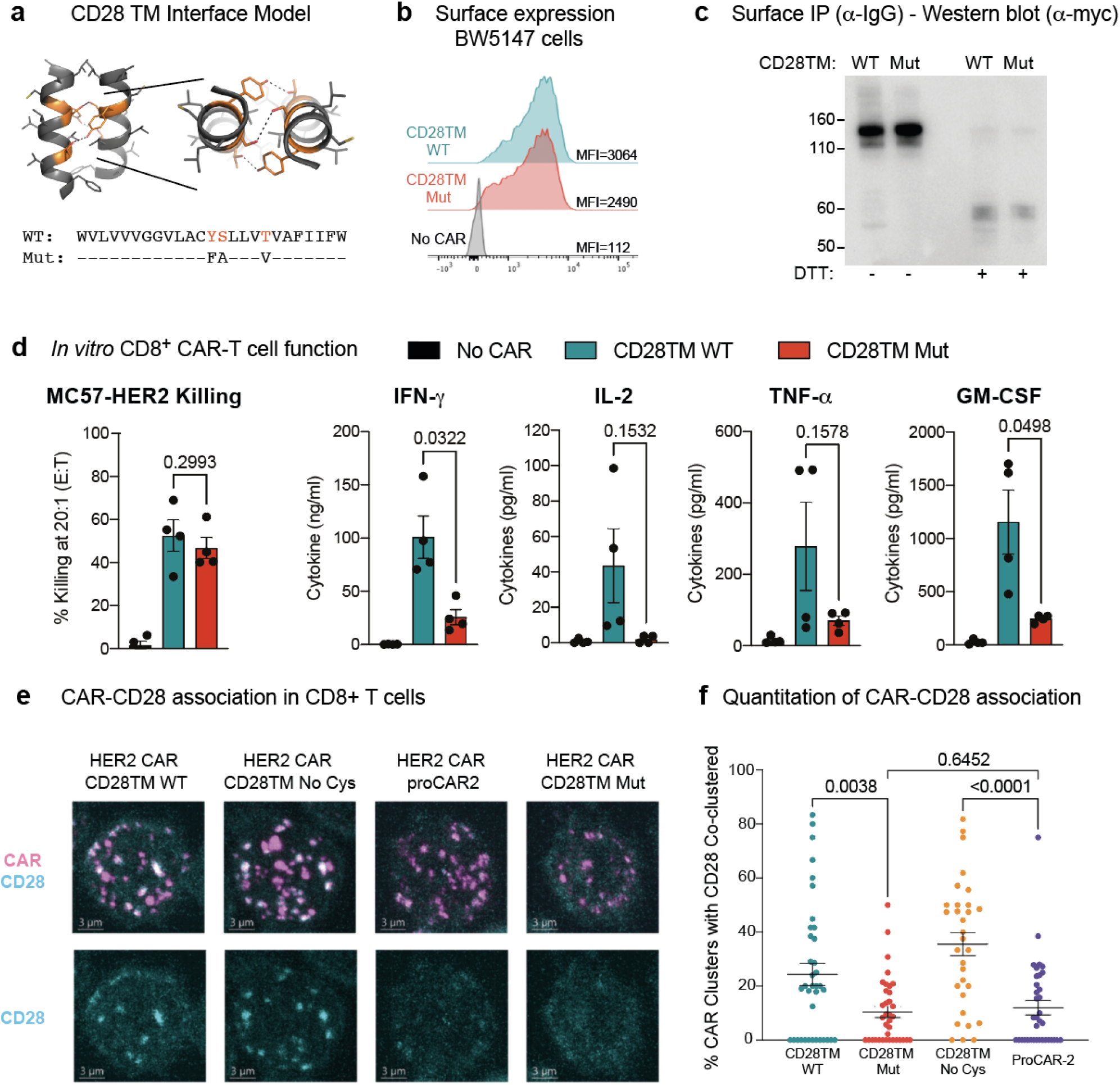
Functional consequences of CAR-CD28 association in CAR T cells. **(a)** Model of the CD28 TM interface generated by mutagenesis of the CD3ζ TMD (PDB:2HAC). Polar residues of the CD28 dimerization motif (orange) with predicted hydrogen bonds depicted (dotted lines). **(b)** Surface expression and **(c)** SDS-PAGE and immunoblot analysis of HER2 CARs possessing WT CD28TM or CD28TM mutations depicted in (a) expressed in the BW5147 cell line. **(d)** Quantitation of target cell killing measured by chromium release assay and cytokine production by primary mouse CD8^+^ CAR T cells in response to the MC57-HER2 target cell line (n=4). Experiments performed as in Figure 3. P-values determined by paired t-tests. **(e)** Representative immunofluorescent confocal images of CAR-CD28 co-clustering in primary mouse CAR T cells. CAR clustering was induced with anti-myc primary followed by crosslinking with fluorescent secondary antibody (magenta). Cells were then labelled for CD28 (cyan). Images are Z-projections over 12**μ**m, scale bar represents 3**μ**m. **(f)** Quantitation of CAR-CD28 co-clustering, each dot representing the percentage of CAR clusters in one cell that co-localised with a CD28 cluster. Lines show mean CAR-CD28 co-clustering percentage/per cell ± SEM, n**≥**30 cells. P-values determined by unpaired *t*-tests.

To directly interrogate potential CAR-CD28 associations in primary CD8^+^ T cells, we examined the four CAR constructs we expect to form dimers in a co-clustering experiment by fluorescence microscopy; these dimers included the CD28TM reference and CD28 TM mutant as well as those that ablate the disulphide linkage for comparison (No Cys and proCAR-2). We found that receptors containing the WT CD28 TMD frequently co-clustered endogenous surface CD28, while the CD28 TM mutant and proCAR-2 did so significantly less frequently (**Figure 4e-f**). These experiments clearly link the CD28 TM interaction motif YSxxxT to high cytokine production in CARs that incorporate this sequence and implicate the recruitment of additional co-stimulatory signalling *via* endogenous T cell CD28 as the cause. They further substantiate that the *de novo* designed TMDs are insulated from these specific interactions.

### *In vivo* antitumor potency scales with proCAR oligomeric state

Short-term *in vitro* tumor cell killing assays do not account for variations in proliferation, survival and cytokine activity that are critical for antitumor activity in a living animal. To evaluate the *in vivo* antitumor potential of proCAR T cells as a function of receptor oligomeric state, we engrafted NOD-SCID-IL2RG^-/-^ (NSG) mice with the aggressive MC38 mouse colon adenocarcinoma cell line engineered to stably express HER2 and treated them one day later with a single intravenous injection of CD8^+^ CAR T cells (**Figure 5a**). Tumors in mice that received empty vector-transduced T cells grew to ethical endpoint (1000 mm^3^) within 14 days, while mice that received proCAR-1, -2 and -3 T cells slowed tumor growth with potency that increased with oligomeric state (**Figure 5b**). ProCAR-3 provided control that most closely resembled the CD28TM reference CAR T cells. The CD28 TM mutant CAR tracked with proCAR-2 and proCAR-3 (**Figure 5c**), confirming that mutation of the YSxxxT association motif recapitulated the general proCAR functional profile *in vivo* as well as *in vitro*. Analysis of mean tumor size at day 14 post-tumor-inoculation (the last day all mice were alive) shows a strong inverse correlation with proCAR oligomeric state (**Figure 5d**). These data show for the first time that, all other features being equal, the potency of antitumor CAR T cell activity scales directly with the oligomeric state of the engineered receptor.

**Figure 5.**
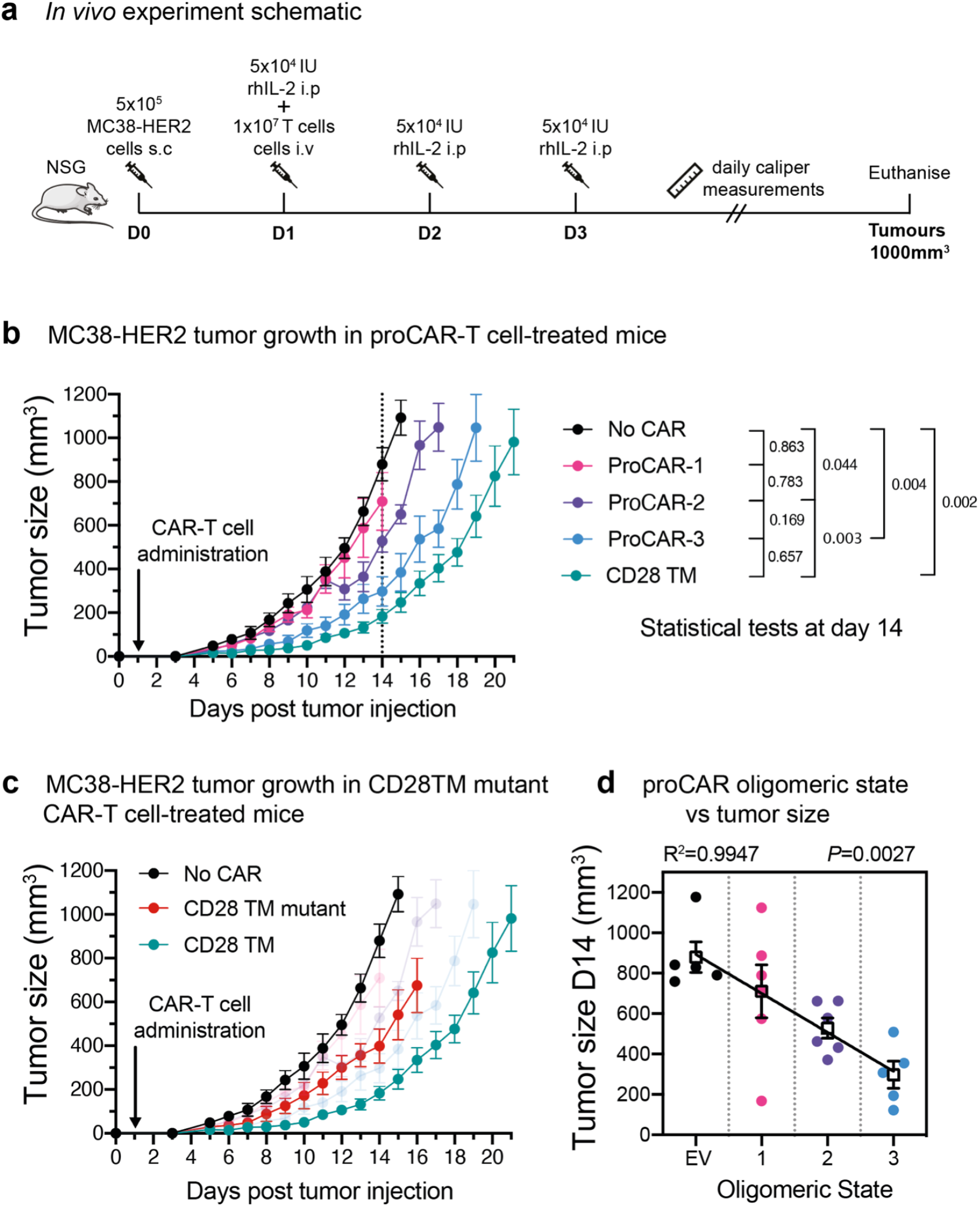
*In vivo* antitumor potency scales directly with proCAR oligomeric state. **(a)** Treatment schedule and experimental setup. NOD-SCID-IL2RG^-/-^ (NSG) mice were injected subcutaneously with MC38-HER2 tumor cells and treated the following day with CD8^+^ T cells delivered intravenously. Mice were supplemented with daily intraperitoneal injections of recombinant human IL-2 from days 1-3, and tumors measured daily until they reached ethical limits. **(b)** Tumor growth over time for No CAR (empty vector), CD28TM WT and proCAR T cell groups (n=5-6 mice/group). Data points represent mean ± error bars showing SEM. Statistical analysis performed using a 2-way ANOVA at day 14. **(c)** Tumor growth over time of CD28TM mutant group superimposed on (b). **(d)** Linear correlation of tumor size on day 14 from (b) vs. proCAR oligomeric state, where the “0” point is provided by empty vector (EV)-transduced T cells. Individual data points are coloured, mean values in white box and error bars indicate SEM. *P*-values indicate the confidence that the slope of the linear regression is non-zero.

### Tetrameric proCAR-4 matches CD28TM CAR tumor control *in vivo* with substantially lower cytokine release *in vitro*

This striking correlation between receptor oligomeric state and functional potency prompted us to push the limits further by designing a tetrameric proMP (proMP C4.1), which features extensive alanine-based complementary packing (**Figure 6a**). The free proMP C4.1 peptide migrates on SDS-PAGE predominantly as a single species at a position indicative of a tetramer (**Figure 6b**), consistent with the observation that complementary apolar packing alone can drive stable MP assembly (Mravic et al., 2019). HER2 proCAR-4 containing the tetrameric proMP C4.1 TMD sequence was well expressed at the surface of freshly isolated mouse CD8^+^ T cells (**Figure 6c** and **Figure S8c**) and supported strong tumor cell killing *in vitro* (**Figure 6d**). This live-cell imaging assay at low effector:target ratio confirmed that oligomeric proCAR T cells and the T cells expressing the reference CD28TM CAR were all potent killers *in vitro*, but the monomeric proCAR-1 T cells clearly segregated with weaker killing. *In vivo*, proCAR-4 T cells displayed a level of MC38-HER2 tumour control that was indistinguishable from the CD28TM reference CAR-T cells (**Figure 6e**), thereby closing the functional gap that was apparent between proCAR-3 and the CD28TM CAR in the previous experiment.

**Figure 6.**
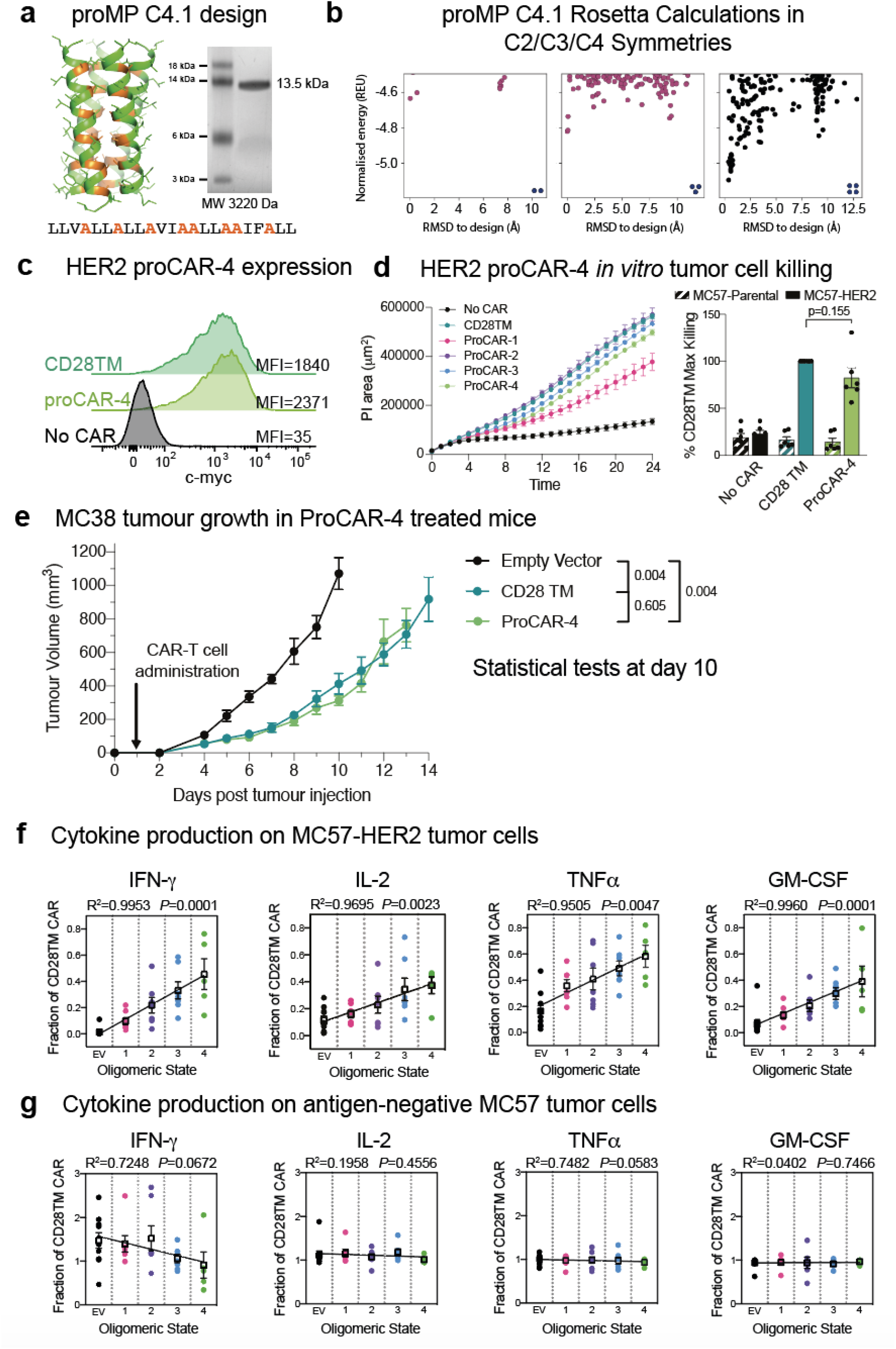
*In vitro* cytokine production scales with proCAR oligomeric state including tetramers. **(a)** SDS-PAGE migration of proMP C4.1 is consistent with a tetramer. Design model and peptide sequence shown for reference. **(b)** Rosetta *ab initio* structure prediction calculations predict that proMP C4.1 preferentially forms a tetramer. **(c)** CAR surface expression on primary mouse CD8^+^ T cells stably expressing CD28TM and proCAR-4 analysed by c-myc staining on flow cytometry. HER2 proCAR-4 was designed using the proMP C4.1 sequence without the final C-terminal leucine as a TMD, inserted as shown in Figure 3a. **(d)** Incucyte killing assay over 24hrs of no CAR, CD28TM and proCAR1-4 T cells on MC57-HER2 target cells at 1:1 effector to target ratio. Comparison of maximum killing for n=6 independent experiments shown between CD28TM vs. ProCAR-4. Data points represent individual experiments, with mean ± SEM error bars plotted. **(e)** Tumor growth over time using the same experimental design in Figure 5a for No CAR (empty vector), CD28TM WT and proCAR-4 T cell groups (n=5-6 mice/group). Data points represent mean ± error bars showing SEM. Statistical analysis performed using a 2-way ANOVA at day 10. **(f-g)** Linear correlation of proCAR oligomeric state vs. IFNγ, IL-2, TNFα and GM-CSF cytokine production (normalized to CD28TM reference) from 24hr co-culture with (e) MC57-HER2 and (f) antigen-negative MC57 tumor cells. Individual data points are coloured, mean values in white box and error bars indicate SEM.

Despite this functional equivalence *in vivo*, proCAR-4 T cells still released significantly lower levels of cytokines than the CD28TM reference CAR T cells *in vitro* (**Figure S10**). However, the tetrameric design trended towards higher levels of all cytokines than the other proCARs. When we normalised cytokine release to the CD28TM reference across all experiments for all proCAR T cells, the combined data revealed strong linear correlations with receptor oligomeric state for all cytokines tested (**Figure 6f**), reflecting a similar relationship to that identified in the *in vivo* tumor control data. Taken together, our results reveal that the high cytokine release stimulated by the CD28TM CAR is largely determined by recruiting native CD28 through the TMD. Yet, amongst the proCAR designs that all eliminate this unintended interaction and thereby reduce cytokine release, the relative cytokine levels scale directly with the receptors’ oligomeric state. This is consistent with a sensitivity to the number of CAR-encoded CD28 and CD3ζ tail sequences that can be engaged by a single antigen-binding event. As expected, cytokine production in response to HER2-negative tumor cells was very low in all constructs (**Figure 6g**), showing that pre-assembled higher-order oligomers did not cause spontaneous antigen-independent activation of cytokine production and still required stimulation. These data confirm a robust linear correlation between CAR oligomeric state and CAR T cell functional output, both *in vivo* and *in vitro*, that extends at least to the tetrameric state.

## Discussion

This work establishes new *de novo* TMD design principles that have direct applications in synthetic biology. Starting from a general methodology for the *de novo* design of membrane-spanning homodimers, we learned that the lowest-energy designed structures systematically exhibited features that are related to protein misfolding, such as self-assembly through multiple alternative interfaces. Furthermore, biochemical and structural analysis noted a surprising tendency of the designs to self-assemble into higher-order oligomers. To counter these unexpected problems, we developed a new strategy that incorporated negative-design principles into an automated design workflow and generated highly expressed and atomically accurate single-span oligomers of defined order. This paved the way to apply *de novo* TMD design to the rapidly developing field of engineered receptors, shedding new light on fundamental structure-function relationships in engineered immune receptors.

The outcomes of the proCAR design experiments revealed two specific mechanistic insights into engineered immune receptor function. First, our results highlight how using natural TMDs can confound predictability and control by encoding unexpected functions. In the HER2 CAR used here, CD28 costimulation is explicitly encoded through the CD28 signaling tail incorporated into the CAR protein but is also amplified through a specific sequence signature in the CD28 TMD that recruits endogenous CD28 into activated CAR complexes. This association likely explains a large portion of the enhanced potency and higher toxicity of CD28 TMD-containing CARs compared to those that use TMDs from CD8 or other proteins (Brudno et al., 2020; Cappell and Kochenderfer, 2021; Davey et al., 2020; Fujiwara et al., 2020; Majzner et al., 2020) and underscores the importance of fully understanding the structure-function relationships in natural TMDs when repurposing them for receptor engineering.

The second major mechanistic insight from this study is that CAR T cell functional potency scales directly with the immune receptor’s oligomeric state when all other features are equal. Systematic and robust interrogation of this relationship has never before been possible because type I single-spanning TMDs with well-characterized oligomeric structures are limited (Trenker et al., 2016) and functional outcomes that depend strictly on oligomeric state are not easily separated from other features and functions of TMDs (Bridgeman et al., 2010, 2014; Wan et al., 2020). Our *de novo* designed proMPs provided a panel of well-characterised, orthogonal TMDs that enabled this finding. The *in vitro* cytokine production and *in vivo* tumor control experiments reported here revealed a striking linear correlation between proCAR oligomeric state and the magnitude of T cell responses.

The ability to broadly attenuate CAR T cell cytokine release while providing a predictable range of functional potencies may have important implications for the development of future cellular immunotherapies. The most effective CAR T cell therapies are accompanied by dangerously high levels of inflammatory cytokine production that cause CRS, which is characterised by fever, hypotension, respiratory distress and multi-organ failure that can be fatal if not carefully managed (Gutierrez et al., 2018; Morgan et al., 2010). The current clinical practice is to manage CRS symptoms with cytokine-blocking antibodies and corticosteroids (Maus et al., 2020; Morris et al., 2021), but approaches to prevent CRS altogether using cytokine gene disruption or modified CAR constructs are areas of active research (Brudno et al., 2020; Sachdeva et al., 2019; Sterner et al., 2019; Ying et al., 2019). The lessons we learned from analysis of monomeric, dimeric and trimeric proCAR designs led to the generation of a tetrameric proCAR with *in vivo* antitumor activity that precisely matched the potent CD28TM design while still providing a 40-60% reduction in inflammatory cytokine release. Other proCAR designs offer greater reductions in cytokine release but also exhibit concomitant loss of antitumor activity *in vivo*. The complete proCAR panel thus provides an opportunity to better balance safety and efficacy in CAR T cell therapies by selecting from a spectrum of designs with defined structure-function relationships. Moreover, TMD modifications do not directly impact either the antigen-binding or signalling domains. This modularity means that proMP TMDs may be easily implemented on the background of any existing single-chain receptor design and can be combined with other modifications in extracellular or intracellular sequences to expand the combinatorial space available for fine-tuning signaling outputs. This flexibility should facilitate screening for an optimal design for each tumor type and target antigen. We anticipate that the proMP design methods and sequences will find additional applications for controlling intermolecular cell-surface protein interactions in a variety of synthetic and biological systems.

## Acknowledgements

We acknowledge the use of the CSIRO Collaborative Crystallisation Centre (C3) for crystallisation screening. X-ray diffraction data were collected at the MX2 beamline of the Australian Synchrotron and we thank the beamline scientists for their technical support. Research supported by a Consolidator’s Grant from the European Research Council (815379) the Dr. Barry Sherman Institute for Medicinal Chemistry and a charitable donation from Sam Switzer and family (to S.J.F.), Australian National Health and Medical Research Council (NHMRC) Project Grant 1158249 (to M.E.C., M.J.C., S.J.F. and M.R.J.) and IRIISS Infrastructure Support (to WEHI), Victorian State Government Operational Infrastructure Support (to WEHI), research grants from the Harry Secomb Foundation and the Percy Baxter Charitable Trust (to M.E.C.) and an equipment grant from the Harold and Cora Brennen Benevolent Trust (to M.E.C.).

## Author Contributions

A.E., J.W., and S.J.F conceived and developed the proMP modelling and design methods and A.E. and J.W. computed the designs. A.E. experimentally tested the designs’ bacterial expression and self-association profiles. N.J.C., J.V.N., R.T. and M.E.C. produced and purified proMP peptides and performed crystallisation screening. N.J.C., J.V.N., R.T. and M.J.C. determined crystal structures. N.J.C. designed and constructed proCARs and performed and analysed all experiments in cell lines using a HER2 CAR construct provided by R.S.C.. A.S.D., N.J.C., M.R.J., M.E.C. and M.J.C designed primary T cell experiments and A.S.D. and N.J.C performed and analysed all experiments in primary T cells. A.S.D. performed *in vivo* experiments. M.E.C. and M.J.C. conceived and supervised proMP peptide production and structure determination strategies and proCAR T cell design and testing strategies. S.J.F. supervised the design process and bacterial expression and self-association experiments. M.E.C. and S.J.F. wrote the manuscript with input from A.E. and M.J.C.. All authors reviewed and edited drafts of the manuscript.

## Competing Interests

A.E., N.J.C., J.W., M.J.C., M.E.C. and S.J.F. are listed as inventors on a patent application covering the proMP sequences and proCARs described herein.

## Supplemental Data

**Figure S1:**
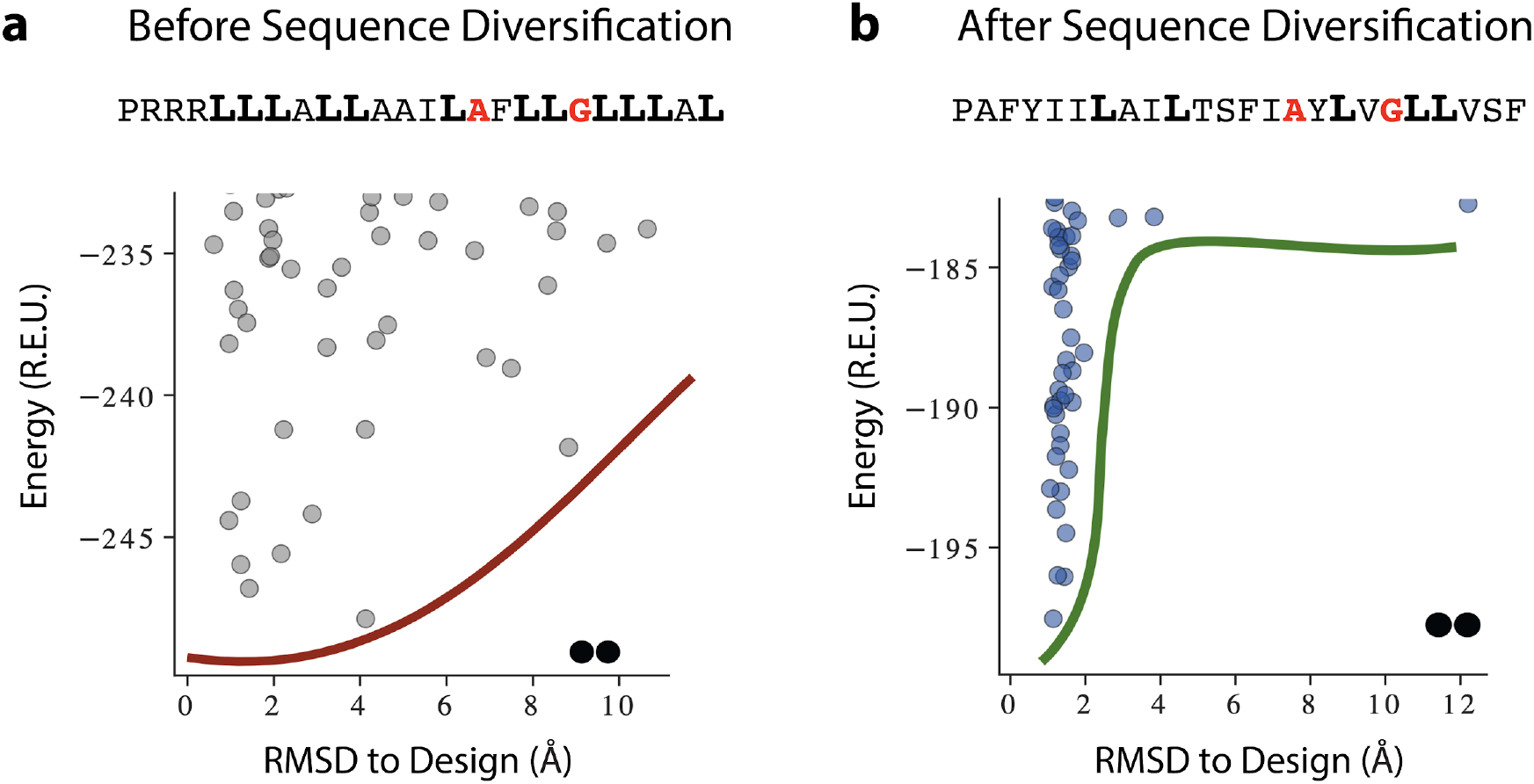
A representative example of proMP sequence diversification resulting in the sequence for proMP 1.8. (**a**) *De novo* designed proMPs exhibited a high proportion of the large, flexible and hydrophobic amino acid Leu (boldface), in accordance with the high lipophilicity of this amino acid according to the dsTβL lipophilicity scale. Forward folding *ab initio* calculations in C2 symmetry, however, exhibited a flat energy landscape with multiple low-energy structures that diverged from the design. (**b**) Simulated annealing Monte Carlo simulations starting from the sequence in (a) augmented with a potential that biased sequence choices to the propensities observed in natural TMDs resulted in sequences with fewer Leu amino acids. Although the Rosetta energies of the designs prior to sequence diversification were more favourable than after (compare the y-axes of the two plots), the sequence-diversified sequences clearly converged to the designed structure. These results suggest that the sequence composition of natural TMDs encodes negative-design principles that ensure folding to a unique conformation. Red one-letter codes indicate positions at the homodimer interface; smooth red and green lines are visual aids.

**Figure S2:**
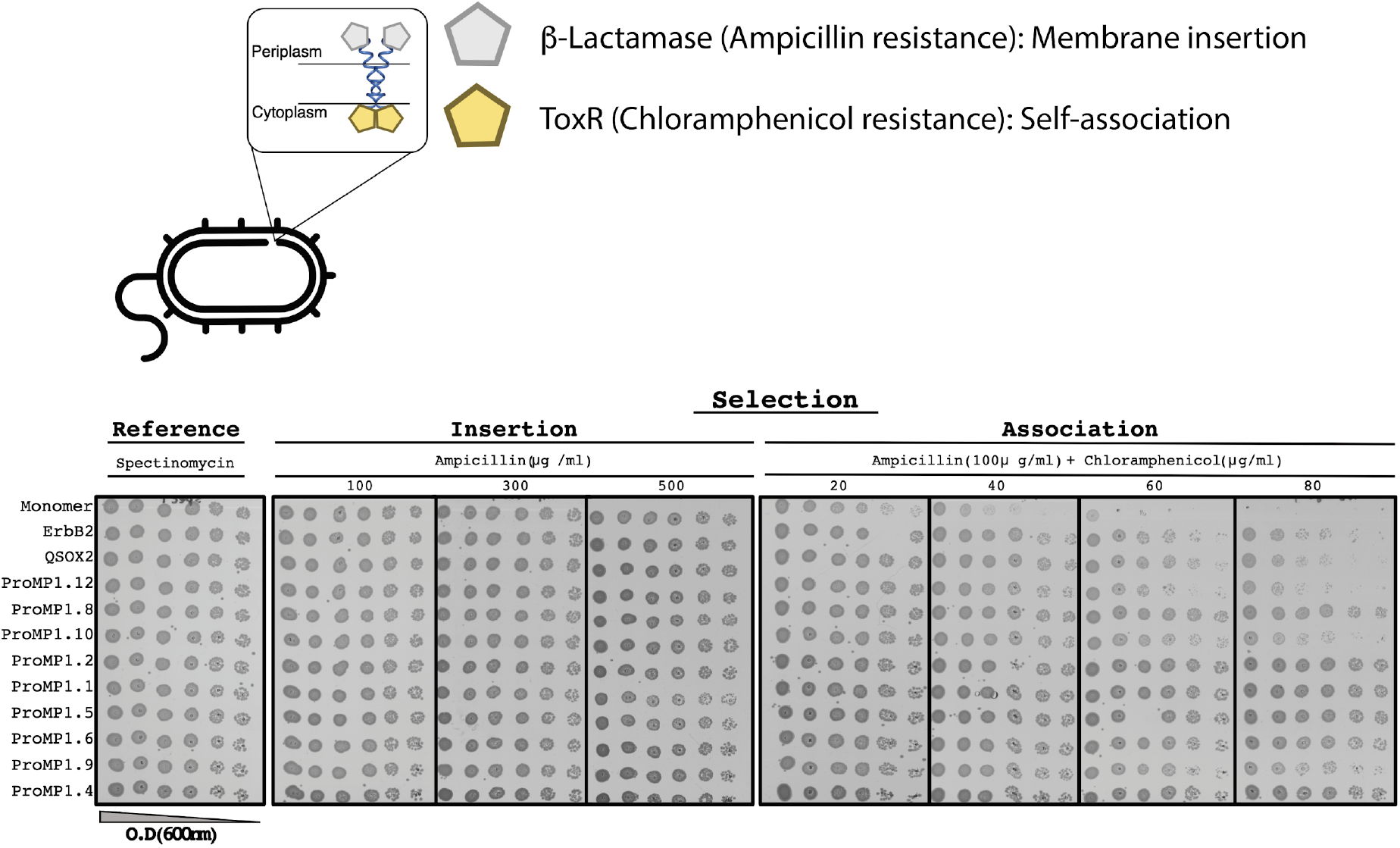
The *E. coli* TOXCAT-β-lactamase (TβL) selection system. (**top**) The construct comprises two selection domains, β lactamase and the transcriptional activator ToxR of the chloramphenicol acetyltransferase. The former is only active in the periplasmic space; the latter depends on dimerisation and is only active in the cytoplasm. Therefore, this system selects bacteria that express a self-associating domain in the inner membrane. (**bottom**) Spectinomycin selects for transformed bacteria; ampicillin selects for membrane insertion; and ampicillin + chloramphenicol selects for membrane insertion and self-association. All the constructs we tested are expressed in the inner membrane, as evidenced by the uniform resistance to ampicillin. The monomeric construct (the C-terminal portion of human L-selectin, CLS) serves as a negative control for self-association and ErbB2 and QSOX2 serve as positive controls based on the TMDs of the respective proteins. Many of the proMPs display a higher level of survival than the positive controls (*e*.*g*., 1.1, 1.2, 1.4, 1.5, 1.6, and 1.8).

**Figure S3:**
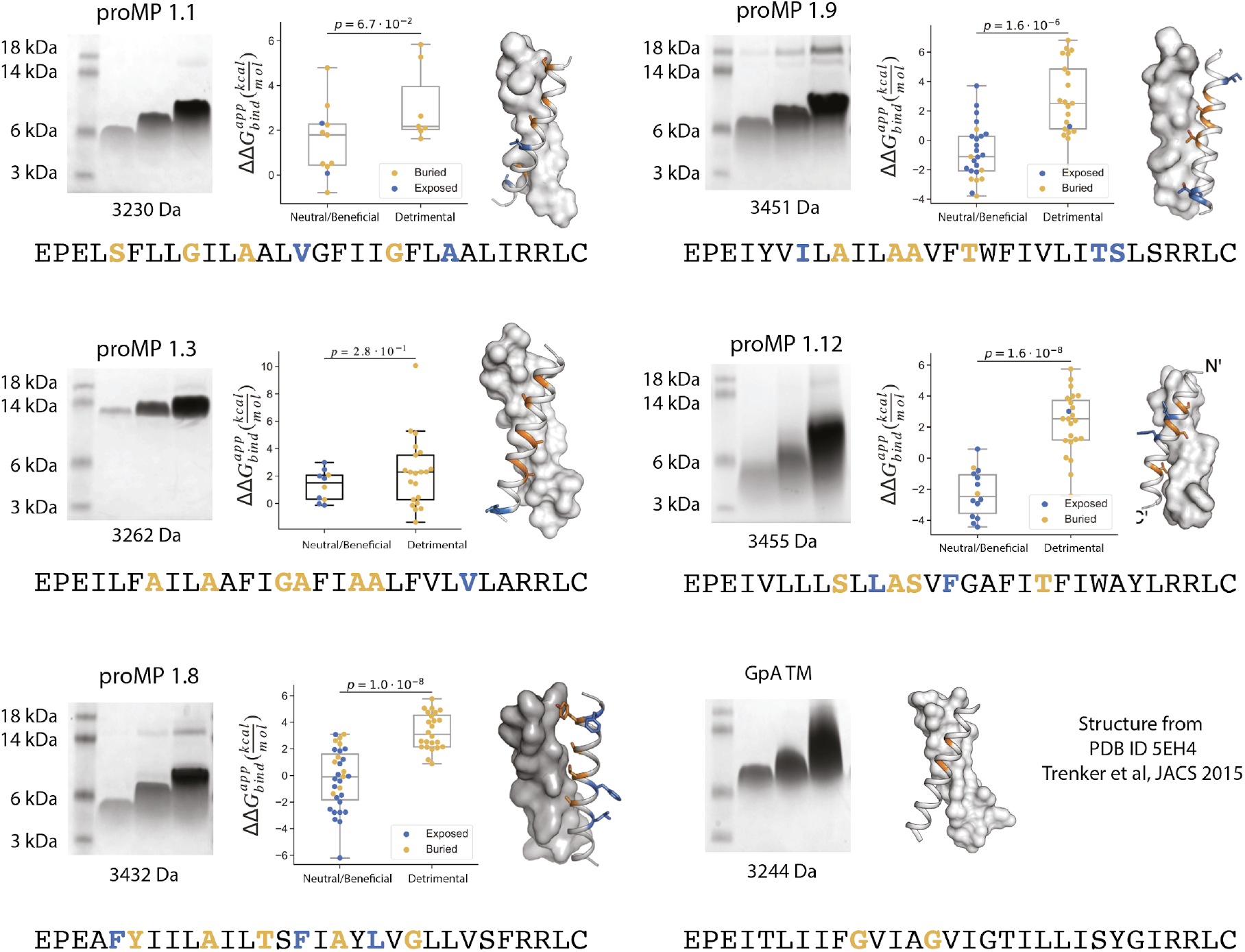
Additional design round 1 sequences, model structures, gel-shift and deep mutational scanning analysis. Each design and a set of single-point mutants were introduced in selected positions (boldface) and assayed using the TβL selection scheme. Amino acids that are positioned at the interface are in orange and those that are lipid-facing are in blue. The mutations were generated using DNA oligos with the degenerate codon NYS, encoding the amino acid identities: Ala, Val, Ile, Leu, Phe, Met, Ser, Thr and Pro (mutations to Pro were excluded from the analysis). We note that proMP 1.3, whose migration on SDS-PAGE indicates an oligomer larger than a dimer, also shows poor segregation of mutations at predicted interface versus lipid-exposed positions in TβL selection, consistent with the formation of a structure that is different from the design model in this case.

**Figure S4:**
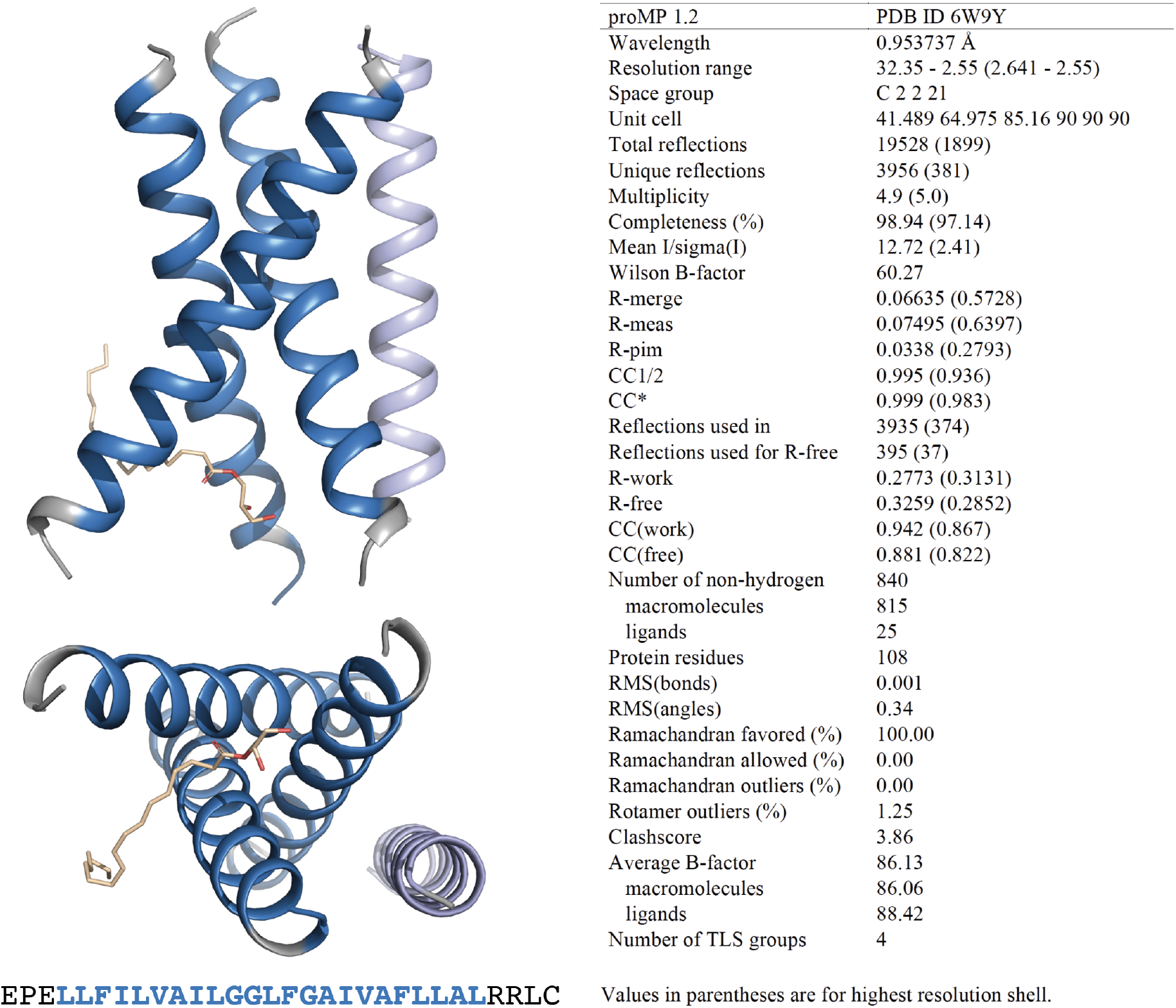
ProMP 1.2 asymmetric unit and structure statistics. Crystals of proMP 1.2 were obtained via LCP crystallisation at 35 mg/ml peptide concentration as described in experimental methods. Screening was performed by the CSIRO Collaborative Crystallisation Centre (C3). Oblong hexagonal discs grown in 25% w/v polyethylene glycol 1500, 10% v/v succinate-phosphate-glycine (pH 6.0) were harvested and frozen in liquid nitrogen using 30% glycerol in precipitant solution as cryo-protectant. ProMP 1.2 packs in the crystal as a trimer (blue) with one monoolein molecule (wheat stick representation) and a less well-ordered helix that is antiparallel in orientation with respect to the trimer (light blue) in the asymmetric unit. Blue ribbons and text represent the designed sequence; gray ribbons and text represent the appended N- and C-terminal sequences included to aid production and crystallisation.

**Figure S5:**
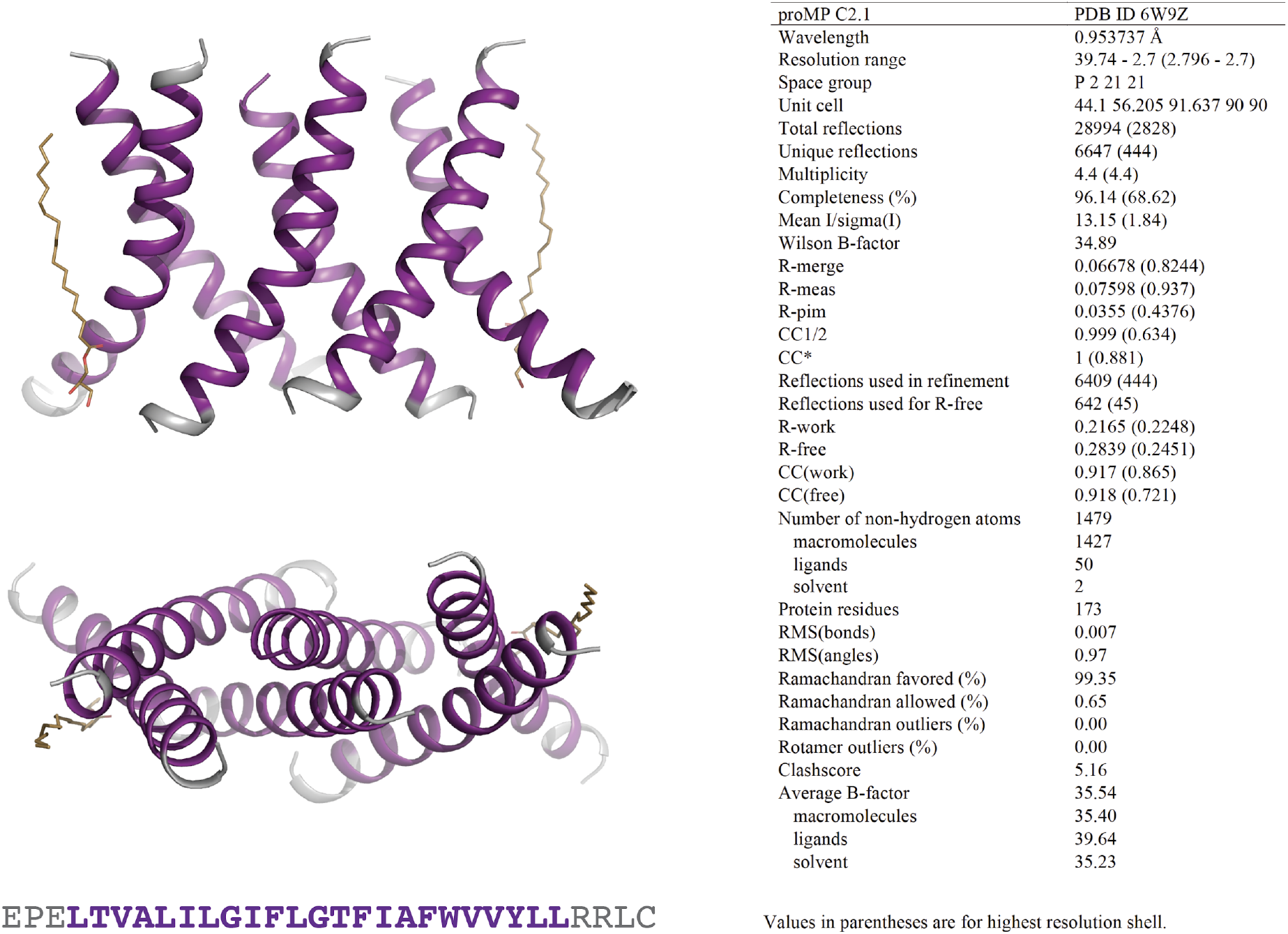
ProMP C2.1 asymmetric unit and structure statistics. Crystals of proMP C2.1 were obtained via LCP crystallisation at 40 mg/ml peptide concentration as described in experimental methods. Screening was performed by the CSIRO Collaborative Crystallisation Centre (C3). Large rhomboid plates grown in 8% v/v 2-methyl-2,4-pentanediol, 0.1 M ADA (pH 6.7), 0.4 M potassium nitrate, 0.1 M tripotassium citrate were harvested and frozen in liquid nitrogen using 30% glycerol in precipitant solution as cryo-protectant. ProMP C2.1 packs in the crystal as a trimer of dimers with two monoolein molecules (wheat stick representation) in the asymmetric unit. Purple ribbons and text represent the designed sequence; grey ribbons and text represent the appended N- and C-terminal sequences included to aid production and crystallisation.

**Figure S6:**
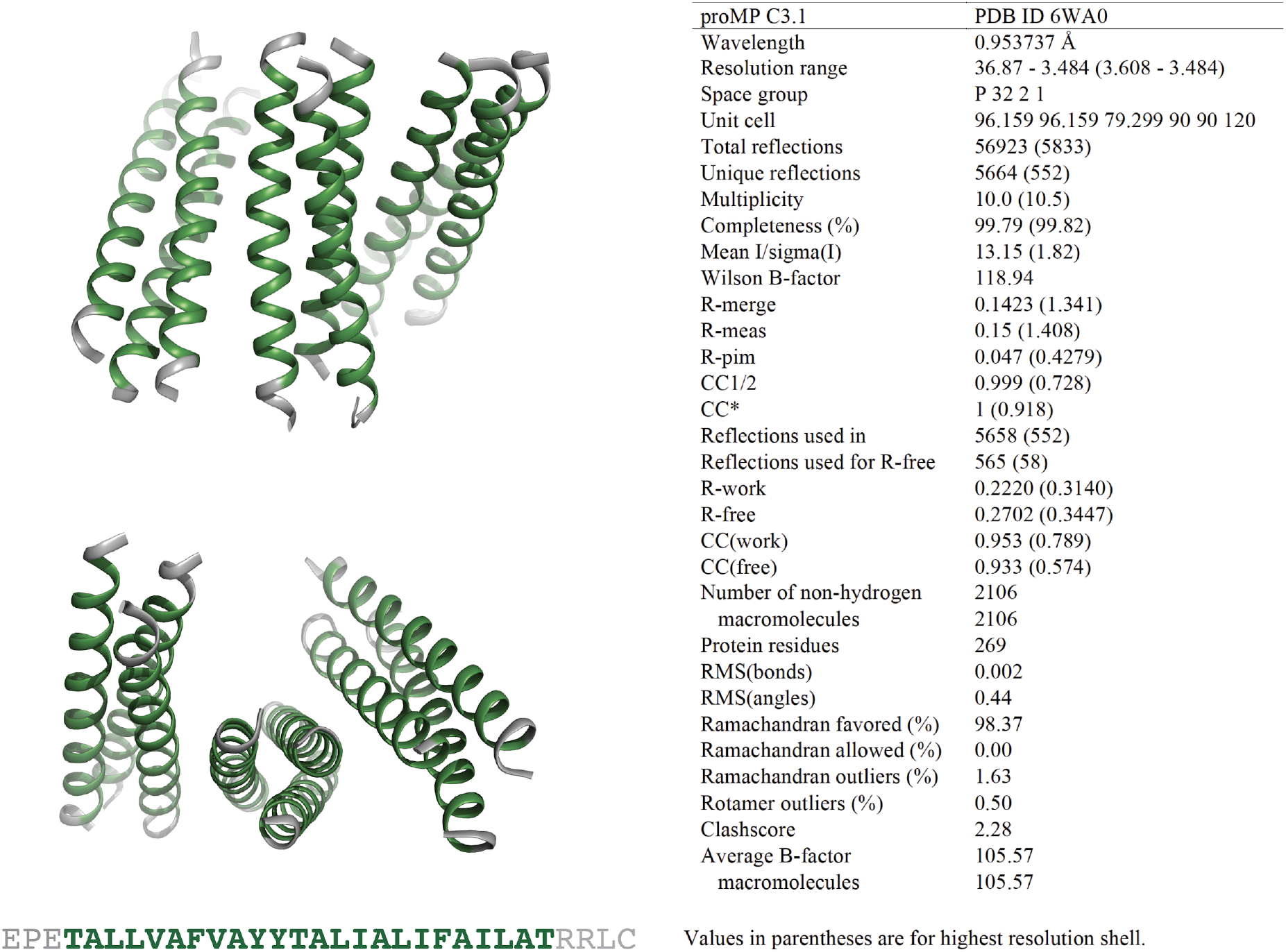
ProMP C3.1 asymmetric unit and structure statistics. Crystals of proMP C3.1 were obtained via sitting-drop vapour diffusion crystallisation at 10 mg/ml peptide in 30 mM C_8_E_4_ as described in experimental methods. Screening was performed by the CSIRO Collaborative Crystallisation Centre (C3). Small ellipsoid crystals grown in 65% v/v 2-methyl-2,4-pentanediol, 0.1 M tris chloride pH 8.0 were harvested and frozen in liquid nitrogen with no additional cryo-protectant. ProMP C3.1 packs in the crystal with a trimer of trimers in the asymmetric unit. Green ribbons and text represent the designed sequence; gray ribbons and text represent the appended N- and C-terminal sequences included to aid production and crystallisation.

**Figure S7:**
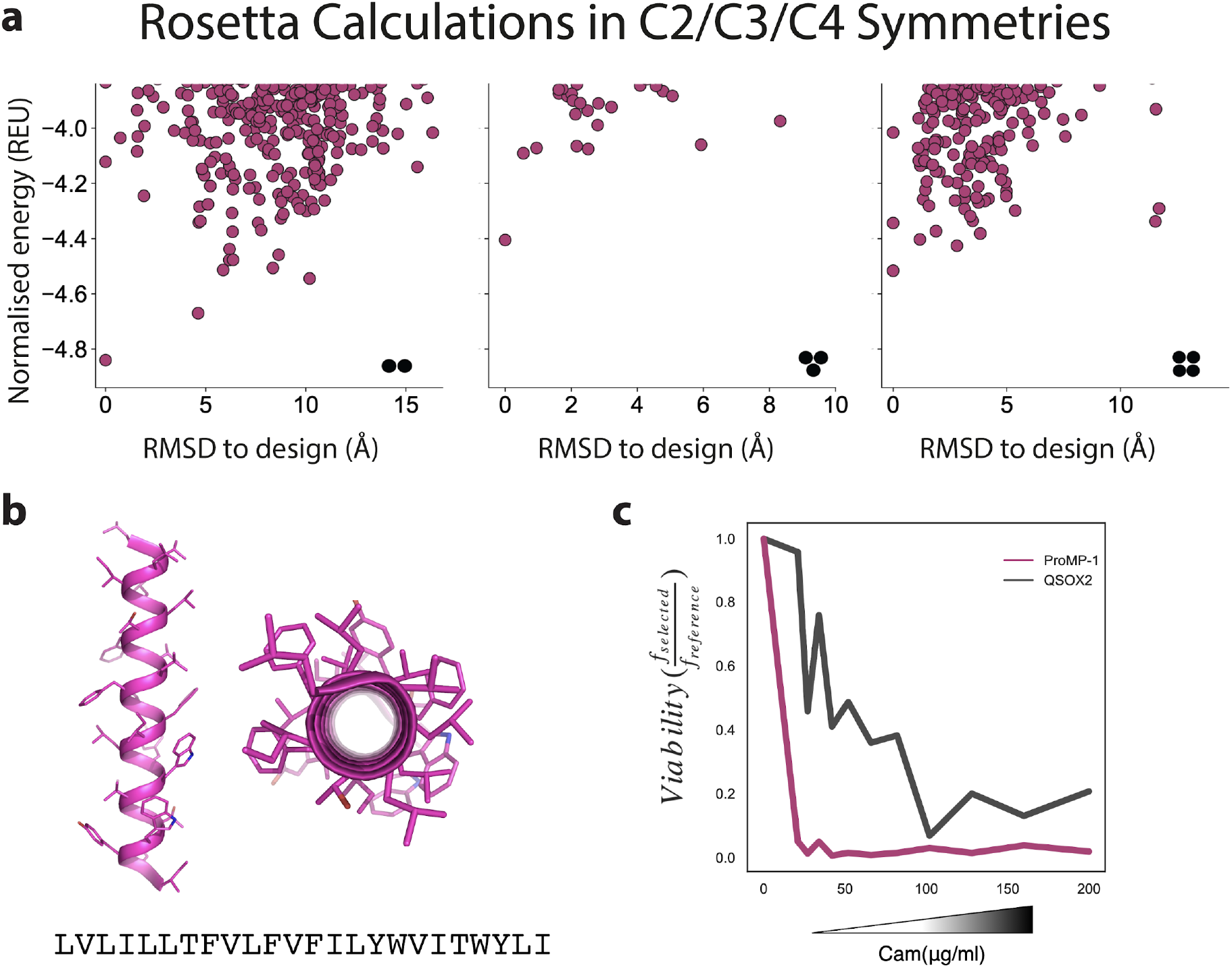
Design of a highly expressed monomeric proMP. (**a**) *Ab initio* structure predictions for proMP-1 in C2, C3 and C4 symmetries exhibit flat energy landscapes suggesting that the design would not form homo-oligomers. (**b**) The design model of proMP-1 shows no flat surfaces that are prone to self-association; yet, the sequence is highly hydrophobic with apparent insertion energy computed using the dsTβL scale of -18.7 kcal/mol (compared to -5.9 kcal/mol for a reference monomer [the human CLS sequence]). (**c**) The design is extremely sensitive to chloramphenicol in the dsTβL assay (compared to the weak natural homodimer QSOX2 which served as a reference), verifying that it does not form homo-oligomers.

**Figure S8:**
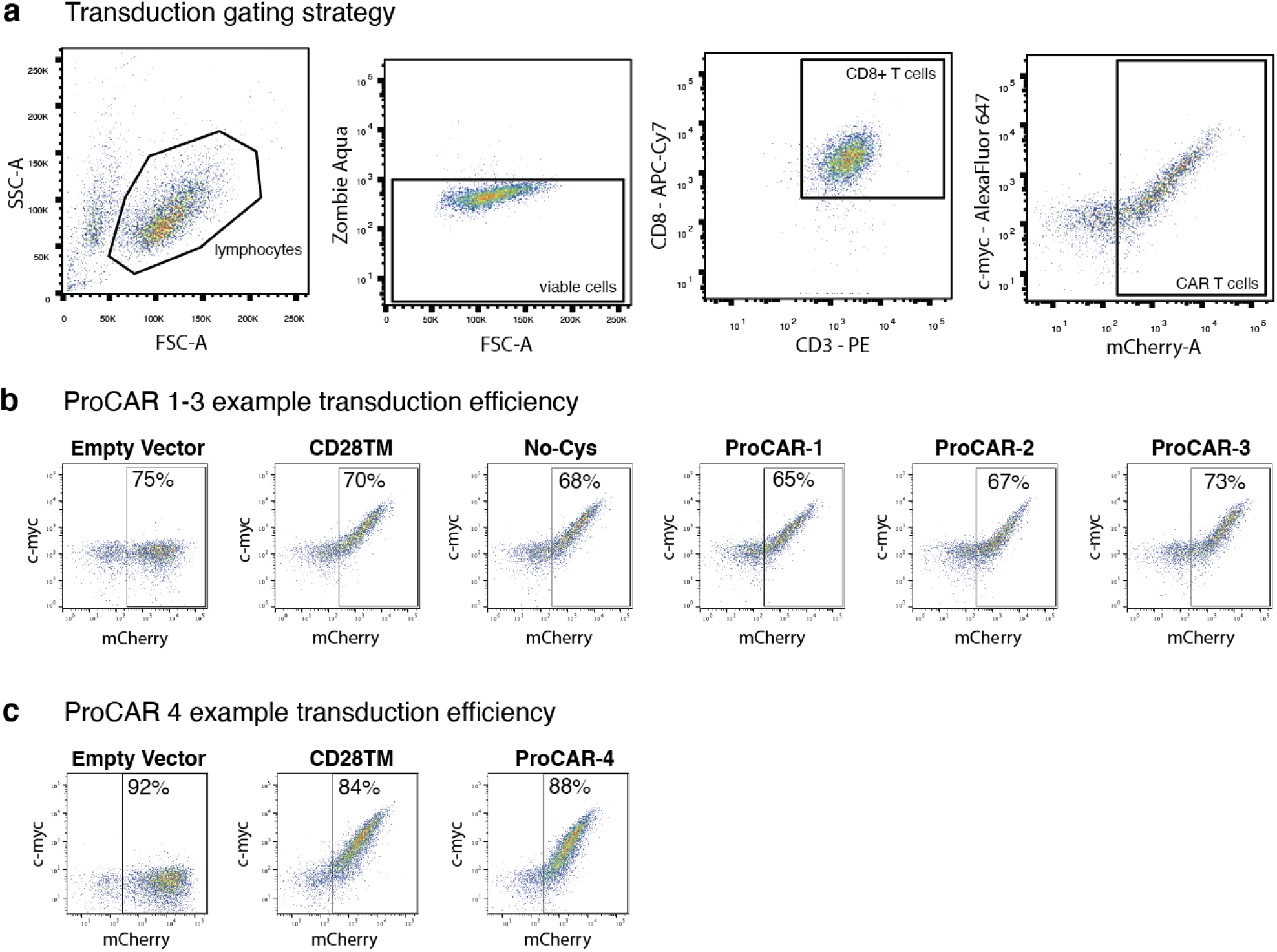
Mouse CAR T cell gating strategy and example ProCAR transduction. (**a**) Flow cytometry gating strategy to determine the transduction efficiency of primary murine CAR T cells. Lymphocytes selected via morphology, live cells selected as zombie aqua negative, T cells selected as CD3^+^CD8^+^ and mCherry^+^ cells defined as CAR T cells. C-myc co-expression with mCherry indicates surface CAR expression. (**b-c**) Example 2D plots showing extracellular c-myc labelling (y-axis) vs. intracellular mCherry (x-axis) of CD3^+^CD8^+^ T cells on day 5 post-transduction with CD28TM CARs and (b) ProCARs 1-3 or (c) ProCAR-4, demonstrating the percentage of cells expressing the CARs. Empty mCherry vector included as c-myc negative control.

**Figure S9:**
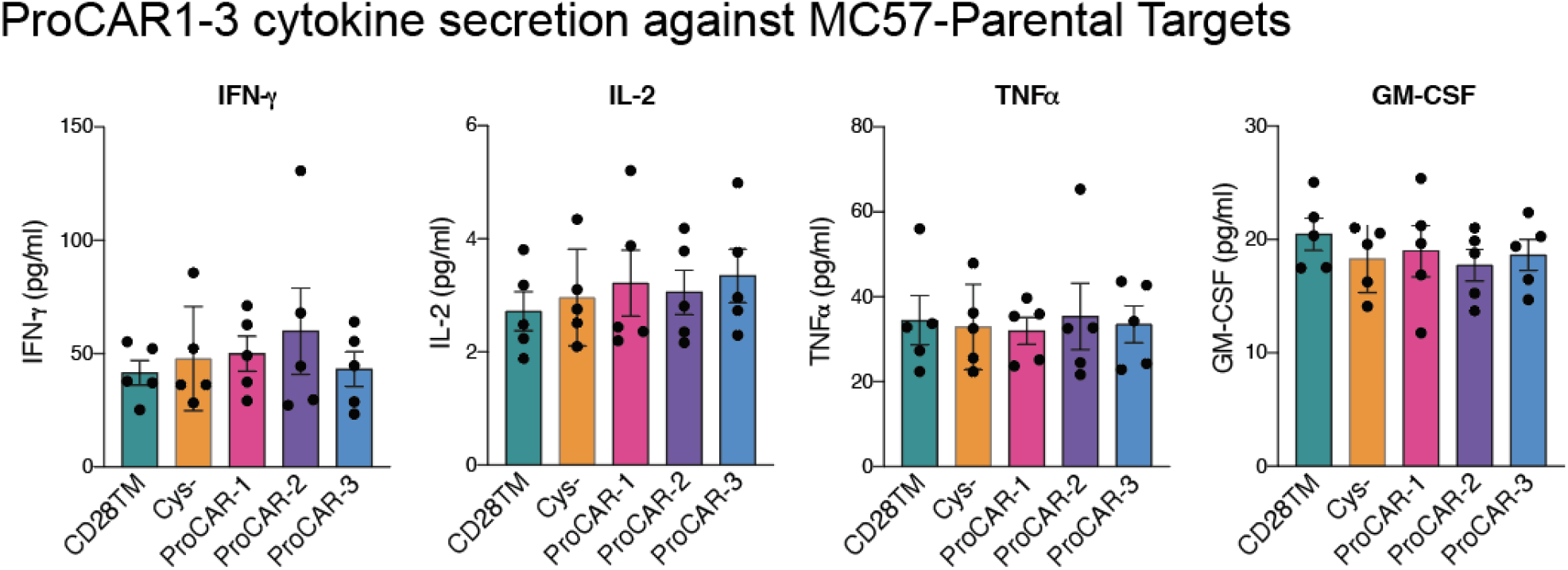
Raw cytokine secretion against MC57-Parental targets for ProCARs 1-3. Cell culture supernatants from CD28TM, no cysteine, proCAR-1, proCAR-2 and proCAR-3 T cells co-cultured with antigen-negative MC57 tumor cells for 24hrs were assessed for cytokine secretion using LEGENDPlex cytokine kits for IFN-γ, IL-2, TNFa and GM-CSF. Mean ± SEM of n=5 independent experiments shown.

**Figure S10:**
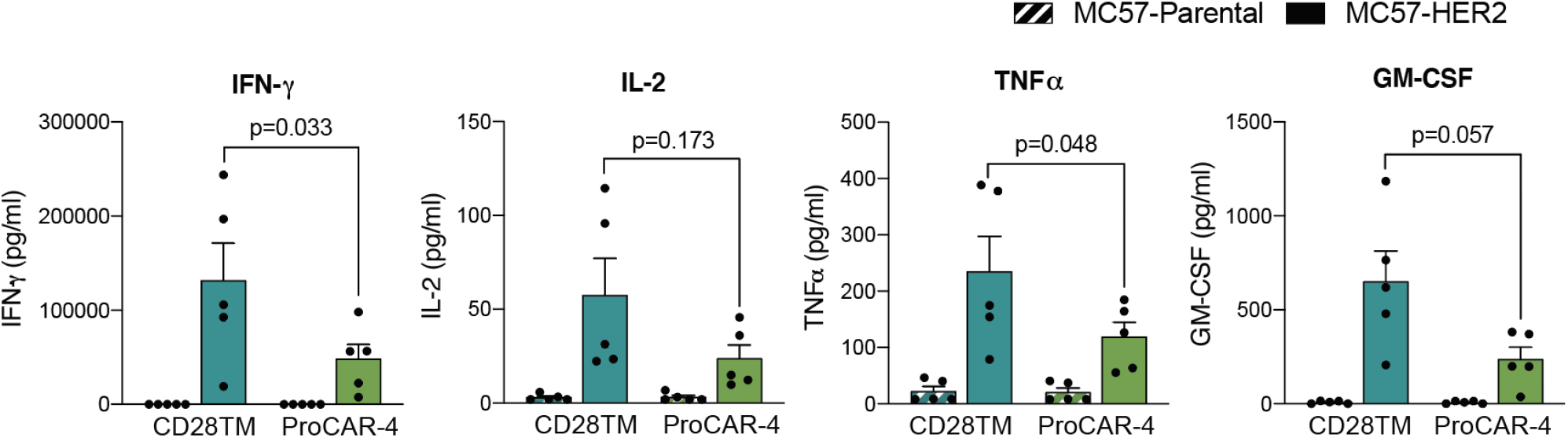
Raw cytokine secretion ProCAR-4 vs. CD28TM. ProCAR-4 and CD28TM CAR T cells were co-cultured with either antigen-negative (MC57-Parental) or MC57-HER2 target tumor cells and incubated for 24hrs. Cell culture supernatants were subsequently collected analysed for cytokine secretion using LEGENDPlex cytokine kits for IFN-γ, IL-2, TNFa and GM-CSF. Mean ± SEM of 5 biological replicates shown.

## Notes

### Summary of Updates

Adding an author that was accidentally missing from the authors list (but was present in the PDF)

## References

Alabanza, L., Pegues, M., Geldres, C., Shi, V., Wiltzius, J.J.W., Sievers, S.A., Yang, S., and Kochenderfer, J.N. (2017). Function of Novel Anti-CD19 Chimeric Antigen Receptors with Human Variable Regions Is Affected by Hinge and Transmembrane Domains. Mol. Ther. 25, 2452–2465.

Arkhipov, A., Shan, Y., Das, R., Endres, N.F., Eastwood, M.P., Wemmer, D.E., Kuriyan, J., and Shaw, D.E. (2013). Architecture and membrane interactions of the EGF receptor. Cell 152, 557–569.

Balakrishnan, A., Rajan, A., Salter, A.I., Kosasih, P.L., Wu, Q., Voutsinas, J., Jensen, M.C., Plückthun, A., and Riddell, S.R. (2019). Multispecific Targeting with Synthetic Ankyrin Repeat Motif Chimeric Antigen Receptors. Clin. Cancer Res. 25, 7506–7516.

Barth, P., and Senes, A. (2016). Toward high-resolution computational design of the structure and function of helical membrane proteins. Nat. Struct. Mol. Biol. 23, 475–480.

Berry, R., and Call, M.E. (2017). Modular Activating Receptors in Innate and Adaptive Immunity. Biochemistry 56, 1383–1402.

Bowie, J.U. (1997). Helix packing in membrane proteins. J. Mol. Biol. 272, 780–789.

Bridgeman, J.S., Hawkins, R.E., Bagley, S., Blaylock, M., Holland, M., and Gilham, D.E. (2010). The optimal antigen response of chimeric antigen receptors harboring the CD3zeta transmembrane domain is dependent upon incorporation of the receptor into the endogenous TCR/CD3 complex. J. Immunol. 184, 6938–6949.

Bridgeman, J.S., Ladell, K., Sheard, V.E., Miners, K., Hawkins, R.E., Price, D.A., and Gilham, D.E. (2014). CD3ζ-based chimeric antigen receptors mediate T cell activation via cis- and trans-signalling mechanisms: implications for optimization of receptor structure for adoptive cell therapy. Clin. Exp. Immunol. 175, 258–267.

Brudno, J.N., Lam, N., Vanasse, D., Shen, Y.-W., Rose, J.J., Rossi, J., Xue, A., Bot, A., Scholler, N., Mikkilineni, L., et al. (2020). Safety and feasibility of anti-CD19 CAR T cells with fully human binding domains in patients with B-cell lymphoma. Nat. Med. 26, 270–280.

Call, M.E., Schnell, J.R., Xu, C., Lutz, R.A., Chou, J.J., and Wucherpfennig, K.W. (2006). The structure of the zetazeta transmembrane dimer reveals features essential for its assembly with the T cell receptor. Cell 127, 355–368.

Cappell, K.M., and Kochenderfer, J.N. (2021). A comparison of chimeric antigen receptors containing CD28 versus 4-1BB costimulatory domains. Nat. Rev. Clin. Oncol.

Cosson, P., Lankford, S.P., Bonifacino, J.S., and Klausner, R.D. (1991). Membrane protein association by potential intrarnembrane charge pairs. Nature 351, 414–416.

Das, R., Andre, I., Shen, Y., Wu, Y., Lemak, A., Bansal, S., Arrowsmith, C.H., Szyperski, T., and Baker, D. (2009). Simultaneous prediction of protein folding and docking at high resolution. Proc. Natl. Acad. Sci. U. S. A. 106, 18978–18983.

Davenport, A.J., Jenkins, M.R., Cross, R.S., Yong, C.S., Prince, H.M., Ritchie, D.S., Trapani, J.A., Kershaw, M.H., Darcy, P.K., and Neeson, P.J. (2015). CAR-T Cells Inflict Sequential Killing of Multiple Tumor Target Cells. Cancer Immunol Res 3, 483–494.

Davenport, A.J., Cross, R.S., Watson, K.A., Liao, Y., Shi, W., Prince, H.M., Beavis, P.A., Trapani, J.A., Kershaw, M.H., Ritchie, D.S., et al. (2018). Chimeric antigen receptor T cells form nonclassical and potent immune synapses driving rapid cytotoxicity. Proc. Natl. Acad. Sci. U. S. A. 115, E2068–E2076.

Davey, A.S., Call, M.E., and Call, M.J. (2020). The Influence of Chimeric Antigen Receptor Structural Domains on Clinical Outcomes and Associated Toxicities. Cancers 13.

Dong, D., Zheng, L., Lin, J., Zhang, B., Zhu, Y., Li, N., Xie, S., Wang, Y., Gao, N., and Huang, Z. (2019). Structural basis of assembly of the human T cell receptor–CD3 complex. Nature 573, 546–552.

Elazar, A., Weinstein, J., Biran, I., Fridman, Y., Bibi, E., and Fleishman, S.J. (2016a). Mutational scanning reveals the determinants of protein insertion and association energetics in the plasma membrane. Elife 5, 12125.

Elazar, A., Weinstein, J.J., Prilusky, J., and Fleishman, S.J. (2016b). Interplay between hydrophobicity and the positive-inside rule in determining membrane-protein topology. Proc. Natl. Acad. Sci. U. S. A. 113, 10340–10345.

Endres, N.F., Das, R., Smith, A.W., Arkhipov, A., Kovacs, E., Huang, Y., Pelton, J.G., Shan, Y., Shaw, D.E., Wemmer, D.E., et al. (2013). Conformational coupling across the plasma membrane in activation of the EGF receptor. Cell 152, 543–556.

Eshhar, Z., Waks, T., Gross, G., and Schindler, D.G. (1993). Specific activation and targeting of cytotoxic lymphocytes through chimeric single chains consisting of antibody-binding domains and the gamma or zeta subunits of the immunoglobulin and T-cell receptors. Proc. Natl. Acad. Sci. U. S. A. 90, 720–724.

Faham, S., Yang, D., Bare, E., Yohannan, S., Whitelegge, J.P., and Bowie, J.U. (2004). Side-chain Contributions to Membrane Protein Structure and Stability. J. Mol. Biol. 335, 297–305.

Feucht, J., Sun, J., Eyquem, J., Ho, Y.-J., Zhao, Z., Leibold, J., Dobrin, A., Cabriolu, A., Hamieh, M., and Sadelain, M. (2019). Calibration of CAR activation potential directs alternative T cell fates and therapeutic potency. Nat. Med. 25, 82–88.

Fleishman, S.J., and Baker, D. (2012). Role of the biomolecular energy gap in protein design, structure, and evolution. Cell 149, 262–273.

Fleishman, S.J., Schlessinger, J., and Ben-Tal, N. (2002). A putative molecular-activation switch in the transmembrane domain of erbB2. Proc. Natl. Acad. Sci. U. S. A. 99, 15937–15940.

Fu, Q., Fu, T.-M., Cruz, A.C., Sengupta, P., Thomas, S.K., Wang, S., Siegel, R.M., Wu, H., and Chou, J.J. (2016). Structural Basis and Functional Role of Intramembrane Trimerization of the Fas/CD95 Death Receptor. Mol. Cell 61, 602–613.

Fujiwara, K., Tsunei, A., Kusabuka, H., Ogaki, E., Tachibana, M., and Okada, N. (2020). Hinge and Transmembrane Domains of Chimeric Antigen Receptor Regulate Receptor Expression and Signaling Threshold. Cells 9.

Gutierrez, C., McEvoy, C., Mead, E., Stephens, R.S., Munshi, L., Detsky, M.E., Pastores, S.M., and Nates, J.L. (2018). Management of the Critically Ill Adult Chimeric Antigen Receptor-T Cell Therapy Patient: A Critical Care Perspective. Crit. Care Med. 46, 1402–1410.

Hartl, F.A., Beck-Garcìa, E., Woessner, N.M., Flachsmann, L.J., Cárdenas, R.M.-H.V., Brandl, S.M., Taromi, S., Fiala, G.J., Morath, A., Mishra, P., et al. (2020). Noncanonical binding of Lck to CD3ε promotes TCR signaling and CAR function. Nat. Immunol. 21, 902–913.

Haynes, N.M., Trapani, J.A., Teng, M.W.L., Jackson, J.T., Cerruti, L., Jane, S.M., Kershaw, M.H., Smyth, M.J., and Darcy, P.K. (2002). Single-chain antigen recognition receptors that costimulate potent rejection of established experimental tumors. Blood 100, 3155–3163.

Hennecke, S., and Cosson, P. (1993). Role of transmembrane domains in assembly and intracellular transport of the CD8 molecule. J. Biol. Chem. 268, 26607–26612.

James, J.R. (2018). Tuning ITAM multiplicity on T cell receptors can control potency and selectivity to ligand density. Science Signaling 11, eaan1088.

Joh, N.H., Wang, T., Bhate, M.P., Acharya, R., Wu, Y., Grabe, M., Hong, M., Grigoryan, G., and DeGrado, W.F. (2014). De novo design of a transmembrane Zn2+-transporting four-helix bundle. Science 346, 1520–1524.

June, C.H., O’Connor, R.S., Kawalekar, O.U., Ghassemi, S., and Milone, M.C. (2018). CAR T cell immunotherapy for human cancer. Science 359, 1361–1365.

Korendovych, I.V., and DeGrado, W.F. (2020). De novo protein design, a retrospective. Q. Rev. Biophys. 53, e3.

Langosch, D., Brosig, B., Kolmar, H., and Fritz, H.J. (1996). Dimerisation of the glycophorin A transmembrane segment in membranes probed with the ToxR transcription activator. J. Mol. Biol. 263, 525–530.

Leddon, S.A., Fettis, M.M., Abramo, K., Kelly, R., Oleksyn, D., and Miller, J. (2020). The CD28 Transmembrane Domain Contains an Essential Dimerization Motif. Front. Immunol. 11, 1519.

Lemmon, M.A., Flanagan, J.M., Hunt, J.F., Adair, B.D., Bormann, B.J., Dempsey, C.E., and Engelman, D.M. (1992). Glycophorin A dimerization is driven by specific interactions between transmembrane alpha-helices. J. Biol. Chem. 267, 7683–7689.

Liu, X., Jiang, S., Fang, C., Yang, S., Olalere, D., Pequignot, E.C., Cogdill, A.P., Li, N., Ramones, M., Granda, B., et al. (2015). Affinity-Tuned ErbB2 or EGFR Chimeric Antigen Receptor T Cells Exhibit an Increased Therapeutic Index against Tumors in Mice. Cancer Res. 75, 3596–3607.

Lu, P., Min, D., DiMaio, F., Wei, K.Y., Vahey, M.D., Boyken, S.E., Chen, Z., Fallas, J.A., Ueda, G., Sheffler, W., et al. (2018). Accurate computational design of multipass transmembrane proteins. Science 359, 1042–1046.

Majzner, R.G., and Mackall, C.L. (2019). Clinical lessons learned from the first leg of the CAR T cell journey. Nat. Med. 25, 1341–1355.

Majzner, R.G., Rietberg, S.P., Sotillo, E., Dong, R., Vachharajani, V.T., Labanieh, L., Myklebust, J.H., Kadapakkam, M., Weber, E.W., Tousley, A.M., et al. (2020). Tuning the Antigen Density Requirement for CAR T Cell Activity. Cancer Discov.

Mata, M., and Gottschalk, S. (2019). Engineering for Success: Approaches to Improve Chimeric Antigen Receptor T Cell Therapy for Solid Tumors. Drugs 79, 401–415.

Matthews, E.E., Zoonens, M., and Engelman, D.M. (2006). Dynamic helix interactions in transmembrane signaling. Cell 127, 447–450.

Maus, M.V., Alexander, S., Bishop, M.R., Brudno, J.N., Callahan, C., Davila, M.L., Diamonte, C., Dietrich, J., Fitzgerald, J.C., Frigault, M.J., et al. (2020). Society for Immunotherapy of Cancer (SITC) clinical practice guideline on immune effector cell-related adverse events. J Immunother Cancer 8.

Morgan, R.A., Yang, J.C., Kitano, M., Dudley, M.E., Laurencot, C.M., and Rosenberg, S.A. (2010). Case report of a serious adverse event following the administration of T cells transduced with a chimeric antigen receptor recognizing ERBB2. Mol. Ther. 18, 843–851.

Morris, E.C., Neelapu, S.S., Giavridis, T., and Sadelain, M. (2021). Cytokine release syndrome and associated neurotoxicity in cancer immunotherapy. Nat. Rev. Immunol.

Mravic, M., Thomaston, J.L., Tucker, M., Solomon, P.E., Liu, L., and DeGrado, W.F. (2019). Packing of apolar side chains enables accurate design of highly stable membrane proteins. Science 363, 1418–1423.

Muller, Y.D., Nguyen, D.P., Ferreira, L.M.R., Ho, P., Raffin, C., Valencia, R.V.B., Congrave-Wilson, Z., Roth, T.L., Eyquem, J., Van Gool, F., et al. (2021). The CD28-Transmembrane Domain Mediates Chimeric Antigen Receptor Heterodimerization With CD28. Front. Immunol. 12, 639818.

Pan, L., Fu, T.-M., Zhao, W., Zhao, L., Chen, W., Qiu, C., Liu, W., Liu, Z., Piai, A., Fu, Q., et al. (2019). Higher-Order Clustering of the Transmembrane Anchor of DR5 Drives Signaling. Cell 176, 1477–1489.e14.

Rafiq, S., Hackett, C.S., and Brentjens, R.J. (2020). Engineering strategies to overcome the current roadblocks in CAR T cell therapy. Nature Reviews Clinical Oncology 17, 147–167.

Sachdeva, M., Duchateau, P., Depil, S., Poirot, L., and Valton, J. (2019). Granulocyte-macrophage colony-stimulating factor inactivation in CAR T-cells prevents monocyte-dependent release of key cytokine release syndrome mediators. J. Biol. Chem. 294, 5430–5437.

Salter, A.I., Pont, M.J., and Riddell, S.R. (2018). Chimeric antigen receptor-modified T cells: CD19 and the road beyond. Blood 131, 2621–2629.

Sterner, R.M., Sakemura, R., Cox, M.J., Yang, N., Khadka, R.H., Forsman, C.L., Hansen, M.J., Jin, F., Ayasoufi, K., Hefazi, M., et al. (2019). GM-CSF inhibition reduces cytokine release syndrome and neuroinflammation but enhances CAR-T cell function in xenografts. Blood 133, 697–709.

Trenker, R., Call, M.J., and Call, M.E. (2016). Progress and prospects for structural studies of transmembrane interactions in single-spanning receptors. Curr. Opin. Struct. Biol. 39, 115–123.

Wan, Z., Shao, X., Ji, X., Dong, L., Wei, J., Xiong, Z., Liu, W., and Qi, H. (2020). Transmembrane domain-mediated Lck association underlies bystander and costimulatory ICOS signaling. Cell. Mol. Immunol. 17, 143–152.

Weinstein, J.Y., Elazar, A., and Fleishman, S.J. (2019). A lipophilicity-based energy function for membrane-protein modelling and design. PLoS Comput. Biol. 15, e1007318.

Wels, W., Harwerth, I.M., Zwickl, M., Hardman, N., Groner, B., and Hynes, N.E. (1992). Construction, bacterial expression and characterization of a bifunctional single-chain antibody-phosphatase fusion protein targeted to the human erbB-2 receptor. Biotechnology 10, 1128–1132.

Woodall, N.B., Yin, Y., and Bowie, J.U. (2015). Dual-topology insertion of a dual-topology membrane protein. Nat. Commun. 6, 8099.

Wu, W., Zhou, Q., Masubuchi, T., Shi, X., Li, H., Xu, X., Huang, M., Meng, L., He, X., Zhu, H., et al. (2020). Multiple Signaling Roles of CD3ε and Its Application in CAR-T Cell Therapy. Cell.

Ying, Z., Huang, X.F., Xiang, X., Liu, Y., Kang, X., Song, Y., Guo, X., Liu, H., Ding, N., Zhang, T., et al. (2019). A safe and potent anti-CD19 CAR T cell therapy. Nat. Med. 25, 947–953.

