## Supplementary material for "*De novo* designed transmembrane domains tune engineered receptor functions": Methods

### Computational methods

Command lines and RosettaScripts(Fleishman et al., 2011) are available below. Rosetta is available at http://www.rosettacommons.org. We used git version b210d6d5a0c21208f4f874f62b2909f926379c0f for all Rosetta calculations.

### Membrane-protein energy function

All atomistic calculations used the Rosetta ref2015_memb energy function(Weinstein et al., 2019). This energy function is based on the recent Rosetta energy function 2015 (ref2015) energetics, which is dominated by van der Waals packing, electrostatics, hydrogen bonding and water solvation, with the difference that in ref2015_memb, the solvation terms are replaced with splines that recapitulate the amino acid based lipophilicity contributions observed in the dsTβL insertion profiles(Elazar et al., 2016). The centroid-level energy function was similarly based on ref2015 with amino acid lipophilicity preferences and a biasing potential that disfavors large interhelical crossing angles that are rarely observed in natural TMDs:

$$penalty = 1.51 \times 10^{-4} \times \theta^{3} - 8.925 \times 10^{-3} \times \theta^{2} + 0.187 \times \theta - 0.532 \quad \mathbf{(1)}$$

Where $\theta$ is the crossing angle between the helix and the membrane normal.

### TMD *de novo* design

3 and 9-mer backbone fragments were generated for a 24-amino acid poly valine extended chain using the Rosetta fragment picker(Gront et al., 2011). The Fold & Dock protocol was used in all design simulations(Das et al., 2009). Briefly, depending on the type of symmetry (C2, C3 or C4), the chains were symmetrically duplicated and each move was applied identically to all chains. Moves included centroid-level fragment insertion and docking, followed by all-atom sequence optimization, and backbone, sidechain and rigid-body minimization. 50,000 independent

trajectories were run and the structure models were filtered using structure and energy-based criteria (the best 1% by system energy, solvent accessible surface area >700 Å; shape complementarity(Lawrence and Colman, 1993) *Sc*>0.6; $\Delta\Delta G_{binding}$ <-15 R.e.u.; helicality <0.1 R.e.u.(Weinstein et al., 2019)). Resulting models were visually inspected and selected for further computational design.

**Sequence diversification**

*De novo* designed sequences exhibited a high propensity of the amino acid Leu. To reduce this bias, we implemented 120 steps of Monte Carlo simulated annealing sequence design. In each step, a random single amino acid change was introduced in any position (mutations were restricted to Gly, Ala, Val, Ile, Leu, Met, Phe, Tyr, or Trp). Following relaxation, the mutant was evaluated on three criteria: $\Delta\Delta G_{binding}$, system energy, and the difference between the amino acid propensities in the design *versus* natural TMDs(Liu et al., 2002) using the following equation($RMSD_{sequence\ comp}$):

$$RMSD_{sequence\ comp} = \sqrt{\frac{\sum_{aa}(f(aa^{design}) - f(aa^{natural}))^2}{L}} \quad (2)$$

Where *f* is the frequency of a given amino acid and *L* is the amino acid sequence length.

The three criteria were then transformed using the “fuzzy”-logic design sigmoidal function(Warszawski et al., 2014):

$$f_x = \frac{1}{1+e^{(x-o)s}} \quad (3)$$

Where *x* is each of the three criteria, and *o* and *s* take the following values: for $\Delta\Delta G_{binding}$ 3 R.e.u. and 1 R.e.u.$^{-1}$, respectively, for system energy 20 R.e.u. and 0.5 R.e.u.$^{-1}$, respectively, and for $RMSD_{sequence\ comp}$ 0.05 and 50, respectively. The *o* thresholds on binding and system energy were computed relative to the energies of the starting model in each design.

The resulting functions were then integrated into a “fuzzy”-logic optimization objective function(Warszawski et al., 2014):

$$f_{\Delta\Delta Gbinding} \wedge f_{system\ energy} \wedge f_{RMSD_{sequence\ comp}} \quad (4)$$

### *Ab initio* structure prediction

Designed sequences were subjected to the membrane fold & dock method essentially as described in (Weinstein et al., 2019). Structure models were filtered using structure and energy-based filters: solvent accessible surface area > 600Å; energy < 0; the distance between the TMD ends along the membrane normal, TMsSpanMembrane > 25Å; fractional agreement between the desired topology for each position (cytosolic, membrane, external) and the designed topology SpanTopologyMatchPos > 0.1).

To evaluate whether the *ab initio* structure predictions are funneled, we computed the *Z*-score:

$$Z = \frac{E_{lowest} - \bar{E}}{STD(E)} \quad (5)$$

Where $E_{lowest}$ is the lowest-energy model with an RMSD of less than 2 Å to the original design model and $E$ represents energies of models with an RMSD > 2 Å and less than 50 R.e.u from $E_{lowest}$. A cutoff of $Z > 2.5$ was typically used to determine whether an energy landscape was funneled.

### Rosetta mutational-scanning calculations

In order to characterise the effects of mutations on the designs' binding energy, we conducted computational mutation scanning using the FilterScan protocol in RosettaScripts (see XMLs section below). If the difference in total energy for a mutation was >2.5 R.e.u., the mutation was predicted to be detrimental, otherwise it is defined as neutral/beneficial.

### Experimental methods

### TOXCAT β-lactamase assays

DNA encoding the designs and controls were cloned into the pMAL_dsTβL vector (Elazar et al., 2016) (available at AddGene #73805) using *XhoI* and *SpeI* restriction sites and selected by growth on spectinomycin and ampicillin in standard concentrations. For positive controls, the natural ErbB2 and QSOXS2 TM domains were chosen (representing strong and weak homo-oligomers respectively (Schanzenbach et al., 2017)). The monomeric C-terminal portion of human L-selectin (CLS)(Elazar et al., 2016; Srinivasan et al., 2011) was chosen as a negative control. Resulting plasmids were transformed into *E. coli* cloni cells (Lucigen), plated on agar plates containing 50 µl/ml spectinomycin followed by overnight growth in a 37 ℃ at 200 rpm. Cultures were then inoculated into fresh LB + 50 µl/ml spectinomycin medium to $OD_{600}$ 1 and then plated on petri dishes containing 50 µl/ml spectinomycin, 100 µl/ml ampicillin or 100 µl/ml ampicillin with a range of different chloramphenicol concentrations.

For single-clone growth assays, 2 µl of cultures at OD 0.1 were diluted and plated on square petri dishes containing different chloramphenicol concentrations (**Extended Data Figure 2**).

**Deep sequencing analysis**

A library encoding all of the designed sequences, controls and single-point mutations in defined positions (using NYS codons to encode hydrophobic and small, mildly polar amino acids) was transformed and grown in large 12 cm petri dishes on different chloramphenicol concentrations (0, 60, 80, 100, and 120 µl/ml for data in Figure 1 and extended Figure 3 and 0, 21, 27, 34, 42, 52, 66, 82, 102, 128, 160, and 200 µl/ml for data in extended Figure 7) overnight. Bacteria were harvested and subjected to deep sequencing library preparation and a protocol and analysis as described in(Elazar et al., 2016).

**Deriving changes in free energy of self-association from the deep mutational scanning data**

From the deep-sequencing analysis, we compute the propensity *p* of each mutant *j* at position *i* relative to the wild type as described in(Elazar et al., 2016):

$$p^{i,j} = \frac{count^{i,j}}{count^{wt}} \quad (6)$$

Where *count* is the number of reads for each variant, adding a pseudo-count of 1 if no reads were detected for the wild type. We then obtain selection coefficients *s* by comparing the selected and reference populations:

$$s^{i,j} = \frac{p^{i,j}_{selected}}{p^{i,j}_{ref}} \quad (7)$$

Where the selected population is selected on ampicillin + chloramphenicol plates (selection for insertion and self-association, respectively) and the reference population is selected only on ampicillin plates (insertion only). At each position *i*, the selection coefficients are transformed to changes in free energy of self association from the wild type identity *wt* to the single-point mutation *j* through the Gibbs free-energy equation:

$$\Delta\Delta G_{i,wt\rightarrow j}^{measured} = - RTln(\frac{s^{i,j}}{s^{i,wt}}) \quad (8)$$

Where $R$ is the gas constant, $T$ is the absolute temperature (310K), and $ln$ is the natural logarithm.

In the TOXCAT-β-lactamase construct, bacterial viability on chloramphenicol depends on the activity of the ToxR chloramphenicol acetyltransferase moiety which in turn depends on oligomer concentrations (Elazar et al., 2016; Langosch et al., 1996; Russ and Engelman, 1999). Oligomer concentrations depend on both membrane insertion and self-association energy (Duong et al., 2007; Elazar et al., 2016). Therefore, the energy computed in Eq. 8 comprises contributions from both membrane insertion (doubled in the case of homodimers) and self-association energy. Thus, to extract the self-association energies for each point mutation, the apparent free energy of self association subtracts the apparent contribution from insertion:

$$\Delta\Delta G_{i,wt\rightarrow j}^{app,assoc.} = \Delta\Delta G_{i,wt\rightarrow j}^{measured} - 2\Delta\Delta G_{i,wt\rightarrow j}^{app,ins.} \quad (9)$$

Where the apparent change in free energy of insertion is computed according to the per amino acid, membrane depth-dependent insertion energies derived from the dsTβL assay in ref. (Elazar et al., 2016).

**DNA sequences of designs**

```
>ProMP 1.1
CCTTTATCTTTCCTCTTAGGGATACTAGCTGCGCTGGTGGGGTTCATCATTGGCTTTTTAGCGGCCTTGAT
T
>ProMP 1.2(trimer; used in proCAR-3)
CCTTTGTTATTTATTCTCGTCGCAATACTTGGAGGCTTATTTGGGGCGATTGTTGCATTCCTTTTGGCGTT
A
>ProMP 1.3
CCGATCCTGTTCGCAATACTGGCGGCTTTCATCGGGGCATTTATAGCTGCCCTGTTCGTGCTAGTATTGGC
A
>ProMP 1.4
```

CCCTTTGGAGCTTTACTAGCAATCATAGCATTCGTCGTAGGAATGTTATTCTCAGCATTCGTTTTACTCAT

C

>ProMP 1.5

CCCTTTAGCTTGTTTTTGGGCGTTATAGCCGGCATTATTGCTGCATTCATCGTTTTATTCCTGGCATTACT

A

>ProMP 1.6

CCTTTTTTATCGCTTGTTGGTGCGCTAATCGGGGCTTTCATAGCATTTATCTTGGCTTTGTTCATTTTGGT

T

>ProMP 1.7

CCGATTCTGATCACTTTGGCAATGCTTACGGGAGCAGTGATTGGGGCGATCTCGTCTTTTCTCCTAGTGTA

T

>ProMP 1.8

CCAGCCTTTTATATTATATTGGCAATTCTCACCTCGTTCATAGCCTATTTGGTGGGTCTACTCGTGTCTTT

T

>ProMP 1.9

CCTATTTACGTTATACTAGCCATCTTGGCCGCGGTATTCACTTGGTTCATAGTCCTTATAACTAGCCTGAG

T

>ProMP 1.10

CCTACGGTTACGAGTGCGATTCTTGGCGTGTCATTCGGTACCTTTATTAGCCTCGTAGCTCTGTGGCTTGC

A

>ProMP 1.11

CCAGTGATTGCAATCTTAACTTTTATAGTCCTCACTGCGATTTCGGGAGCGCTGCTCGCTGTTTGGTTCTC

C

>ProMP 1.12

CCCATCGTCTTGCTCCTCAGTCTACTCGCCAGTGTATTTGGGGCGTTCATCACATTTATTTGGGCTTACTT

G

>ProMP C1 (monomer; used in proCAR-1)

CTGGTGCTGATTCTGCTGACCTTTGTGCTGTTTGTGTTTATTCTGTATTGGGTGATTACCTGGTATCTGAT

T

>ProMP C2.1 (dimer; used in proCAR-2)

CCGCTGACCGTGGCGCTGATTCTGGGCATCTTCCTGGGCACCTTTATTGCGTTTTGGGTGGTGTATCTGCT

G

>ProMP C3.1

ACCGCGCTGCTGGTGGCGTTTGTGGCGTATTATACCGCGCTGATTGCGCTGATTTTTGCGATTCTGGCGAC

C

>ProMP C4.1 (tetramer; used in proCAR-4)

```
CCCCTTTTAGTCGCCTTATTGGCGCTGCTTGCTGTAATCGCCGCATTATTAGCAGCTATCTTTGCATTGCT
G
>CLS
CCGCTGTTCATCCCGGTTGCAGTTATGGTTACCGCTTTTAGTGGATTGGGGTTTATCATCTGGCTGGCTAC
>ErbB2
TCTATCATCTCTGCGGTGGTTGGCATTCTGCTGGTCGTGGTCTTGGGCGTGGTCTTTGGCATCCTGAT
>QSOX2
AGCCTATGCGTTGTTTTATACGTGGCATCTAGTTTATTTATGGTCATGTACTTCTTC
```

#### proMP peptide production

Peptides were produced recombinantly as 9His-trpLE fusion proteins in *E. coli* following a previously published protocol(Sharma et al., 2013). To aid purification, analysis and crystallisation, all designed sequences were modified to include Glu-Pro-Glu at the amino terminus and Arg-Arg-Leu-Cys at the carboxy terminus based on the favourable properties of the glycophorin A TMD fragment whose structure has been previously determined by x-ray crystallography(Trenker et al., 2015). Dissolved fusion protein from inclusion bodies was purified on Nickel affinity resin, cyanogen bromide digested and reverse-phase HPLC purified following the published procedure(Sharma et al., 2013) with the following modifications: the C-terminal Cys sulfhydryl group was protected using 10 mM S-methyl methanethiosulfonate (MMTS, Sigma-Aldrich) during lysis and inclusion body solubilisation and peptides were at no time disulfide linked. HPLC-purified peptides were stored as lyophilised products at room temperature until needed.

#### SDS-PAGE analysis

Samples were prepared by drying indicated amounts of each purified peptide taken from dried and weighed product redissolved in 1,1,1,1,1,1-hexafluoroisopropanol (HFIP, Merck). Samples were lyophilised, re-dissolved in 25 µl 1x NuPAGE LDS sample buffer (Thermo Fisher Scientific) and heated for 1 minute at 95°C. Cooled samples were separated on 12% NuPAGE Bis-Tris gels (Thermo Fisher Scientific) at 200V for 40 minutes and visualised by staining with Coomassie Blue R-250 (Bio-Rad).

#### Crystallisation screening and structure determination

#### ProMP crystallisation in LCP

For reconstitution into LCP, lyophilised peptide was weighed and co-dissolved with appropriate amounts of monoolein (Nu-Chek Prep) in HFIP. Solvent was removed under streaming nitrogen, followed by lyophilisation overnight. Peptide-monoolein mix was heated (52°C) until liquid and mixed 3:2 with 10 mM Tris pH 8.0 for LCP formation using coupled 100 µl gastight Hamilton syringes (Formulatrix) at room temperature. For screening, LCP mixture was dispensed in 100 nl drops onto 96-well glass plates (Molecular Dimensions) with 1000 µl of precipitant solution using a Mosquito LCP robot (TTP Labtech) at room temperature. Plates were sealed and kept at 20°C in a Rock Imager 1000 (Formulatrix) for incubation and monitoring of crystal formation.

**ProMP crystallisation in detergent**

For reconstitution with detergent, lyophilised peptide was weighed and dissolved in 30 mM detergent ($C_8E_4$; Anatrace, $C_8E_5$; Anatrace) in HFIP. Solvent was removed under streaming nitrogen followed by lyophilisation overnight. Peptide-detergent mix was reconstituted in 10 mM Tris pH 8.0. For screening, peptide-detergent mixture was dispensed in 150 nl drops onto SD-2 plates (IDEX Corp) with 150 nl of precipitant solution using a Crystal Phoenix robot (Art Robins Instruments) at room temperature. Droplets were equilibrated against 50 nl of crystallant in the reservoir. Plates were sealed and kept at 20°C in a Rock Imager 1000 (Formulatrix) for incubation and monitoring of crystal formation.

**Data Collection and Structure Determination**

Data were collected on the MX2 beamline of the Australian Synchrotron at a wavelength of 0.9537 Å and a temperature of 100 K. Data were indexed and scaled using XDS(Kabsch, 2010) and Aimless(Winn et al., 2011). Structure factor amplitudes were obtained using cTruncate[15]. 6W9Y was solved with Phaser(McCoy et al., 2007) by molecular replacement using the GpA monomer helix as a search model (PDB code 5EH6(Trenker et al., 2015)). 6W9Z was solved with Phaser by molecular replacement using 5EH6 mutated to the proMP C2.1 sequence in *Coot(Emsley et al., 2010).* 6WA0 was solved with Phaser by molecular replacement using the designed trimer as a search model. This resulted in a model that contained good density for two chains, with the final chain of the trimer considerably worse. The third chain was removed and a second molecular replacement job was performed with the first two chains fixed in place and a single helix from the model trimer used as a search model. This resulted in placement of the third helix in an antiparallel direction with respect to the other two chains and this was judged as correct based on comparison of overall Rfree of each model, average B factors of each chain and visual inspection of the electron density in *Coot(Emsley et al., 2010)*. Iterative rounds of refinement and model building were performed in PHENIX(Liebschner et al., 2019) and *Coot*[17].

**ProCAR construct preparation**

The HER2 specific CAR used was based on a previously described construct(Haynes et al., 2002)restriction digest sites removed and were gibson cloned together and inserted into EcoRI/XhoI digested pMSCV-IRES-mCherry-II vector (NEB Gibson Assembly Master Mix Cat# E2611L). The CAR construct contains the FRP5 anti-HER2 scFv, Myc tag, human CD8α stalk, human CD28 TM/tail and human CD3ζ tail sequences. PCR primers were used to generate a cysteine to alanine mutation in the CD8α stalk region to prevent covalent dimerisation. Overlapping PCR was used to generate CARs with altered TM domains on the background of the cysteine-mutated CD8α stalk. These constructs were inserted into the pMSCV-IRES-mCherry-II vector via EcoRI/XhoI restriction sites.

**Surface IP and immunoblot analysis**

$2x10^7$ cells per sample were pelleted and washed twice with phosphate-buffered saline (PBS) prior to coating with 20 µg/ml polyclonal anti-mouse IgG for 45 minutes on ice. Cells were washed twice with PBS and lysed in 200 µl PBS/1% IGEPAL-640/P8340 protease inhibitor/10 mM iodoacetamide for 30 mins on ice. Lysate was centrifuged at 20,000 g for 10 mins, 10 µl of cleared lysate was taken for 5% input controls with remainder being added to 20 µl Thermofisher Protein G agarose beads and rotated in cold room for 2 hrs. Beads were washed with lysis buffer twice then eluted with LDS and boiled. Samples were run on SDS-PAGE and transferred for blotting with 1:2,000 anti-Myc primary antibody (Cell signalling #2276) and 1:20,000 anti-mouse IgG HRP secondary (Sigma Aldrich A0168).

**Animals**

All mice were of an inbred C57B/6J or NOD.Cg-Prkdc$^{scid}$IL2rg$^{tmWjl}$/SzJ (NSG) genetic background. All animal experiments were approved and performed in accordance with the regulatory standards of the Walter and Eliza Hall Institute Animal Ethics Committee (Approval: WEHI-2019.020).

**Mouse CD8$^+$ T cell isolation and culture**

Single-cell suspensions of peripheral lymph nodes from 6-8 week old C57B/6 mice were prepared by mechanically dissociating through a 70 µm cell strainer (BD Biosciences) into cold PBS. CD8$^+$

T cells were subsequently selected using the EasySep™ mouse CD8a positive Kit II (Stem Cell Technologies) according to manufacturer's instructions. Purity was confirmed as >95% using LSR II Fortessa (BD Bioscience), FACSymphony™ (BD Biosciences) or Aurora (Cytek). $CD8^+$ T cells were subsequently activated by incubating overnight with Mouse T-Activator CD3/CD28 Dynabeads™ (Gibco) at a bead to cell ratio of 1:1 in mouse T cell medium (mTCM) consisting of Roswell Park Memorial Institute (RPMI) 1640 Medium (Gibco) supplemented with foetal bovine serum (10%; Bovogen Biological), L-glutamine (2 mM; Gibco), sodium pyruvate (1 mM; Gibco), non-essential amino acids (1x; Sigma-Aldrich), β-mercaptoethanol (50 μM; Sigma-Aldrich) and recombinant human IL-2 (100 IU/ml; PeproTech). Following removal of magnetic beads, T cells were maintained at $1x10^6$ cell/ml in mTCM.

**BW5147 and Primary Mouse CAR-T cell generation**

Retrovirus for all T cells was produced using calcium phosphate transfection of HEK293T cells. BW5147 cells expressing a destabilised-GFP NFkB reporter element were mixed 1:1 with filtered viral supernatant at a final density of 2.5x10^5 cells/ml. Polybrene transfection reagent (Merck) was added to a final concentration of 8 μg/ml polybrene prior to spinfection (2,500 rpm, 37℃, 45 mins). For primary mouse T cells, plates were coated with 32 μg/ml retronectin (Takara Bio) for 24 hrs before plating of $1x10^6$ cells in 1ml viral supernatant and performing a spinfection (2,500 rpm, 37℃, 45 mins). Viral supernatant was removed after 16 hrs and replaced with RPMI supplemented with foetal bovine serum (10%; Bovogen Biological) and L-glutamine (2 mM; Gibco) for BW5147 cells, or mTCM for primary T cells.

**CAR T cell SKBR3 co-culture assay**

$5x10^4$ cells/cell-line were aliquoted onto a confluent layer of SKBR3 cells in a 96-well plate at specified time-points. After 8 hrs, all time points were removed from plate and stained with 1:200 anti-CD69 (Biolegend #104524) on ice for 45 mins. Samples were analysed on a LSR Fortessa X20 (BD Bioscience) and data were analysed using FlowJo™ v10 software.

**Flow Cytometry**

For $CD8^+$ T cell selection and transduction efficiency verification, single cell suspensions were washed and stained with Live/Dead marker Zombie Aqua™ (BioLegend) for 15 mins at RT in PBS, before washing and labelling for at least 30 mins on ice with a panel of monoclonal antibodies (mAbs) including: anti-mouse CD3ε PE (clone 145-2C11, Biolegend), anti-mouse CD8α APC-Cy7 (clone 53-6.7, Biolegend) and anti-mouse Myc-Tag Alexa Fluor® 647 (clone 9B11, Cell Signalling). All samples were analysed with an LSR II Fortessa (BD Bioscience), FACSymphony™ (BD Biosciences) or Aurora (Cytek) and data were analysed using FlowJo™ v10 software.

**Chromium Release Killing Assay**

Standard $^{51}Cr$ release assays were conducted to assess CAR T cell cytotoxicity by measuring release of radioactivity into culture supernatants as cells are lysed. Target MC57 mouse fibrosarcoma cells stably expressing human HER2 (MC57-HER2) were pre-loaded with 100 µCi $^{51}Cr$ for 1 hour at 37°C, washed three times and then $2x10^4$ tumour cells were co-incubated with CAR T cells at effector-to-target (E:T) ratios ranging from 40:1 to 1.25:1. Supernatants were harvested after 4 hrs of co-incubation, plated onto a 96-well scintillator coated LumaPlate (PerkinElmer) and $^{51}Cr$ release quantified using a MicroBeta$^2$ Microplate Counter (PerkinElmer). Target tumour cells incubated in a 5% Triton X-100 solution were used as a maximum release control, while tumour cells incubated in mTCM alone were used as a spontaneous release control. Percent lysis was calculated as: % lysis = ((Experimental release – Spontaneous release) ÷ (Maximum release – Spontaneous release)) 100. Data in Figure 3f and Figure 4d are derived from the 20:1 E:T ratio where killing was maximal for all constructs.

**IncuCyte Killing Assay**

To measure tumour cell death over time, the live-cell imaging system IncuCyte® SX3 or SX5 was used. In this assay, $8x10^3$ target tumour cells per well were plated in a 96-well plate in triplicate, and the following day $CD8^+$ T cells were added at an effector to target ratio of 1:1 in mTCM media. 50 µM propidium iodide (PI; Sigma-Aldrich) was added to each well as a surrogate marker of cell death. Wells were subsequently imaged every hour for 24 hrs, with phase and PI fluorescence recorded. All images were analysed using the IncuCyte® Analysis Software program, where the average PI area (µm) was calculated for each individual well from at least two images per time point. Target tumour cells incubated in a 1.2% (w/v) Saponin (Sigma-Aldrich) solution were used as a positive PI release control while tumour cells incubated in mTCM alone were used as a background PI release control for PI area calculations. Data in figure 6d shows all biological replicates and time points graphed as PI area (y-axis) vs. time (x-axis) using GraphPad Prism v 9.0.0.

**Cytokine Bead Array**

To assess cytokine secretion by CAR T cells, cytokine bead arrays on co-culture supernatants were performed. Murine CAR T cells ($1x10^5$ cells) were washed once in PBS and co-incubated with either mTCM alone, a 1:1 bead to cell ratio of Mouse T-Activator CD3/CD28 Dynabeads™ (Gibco) as a positive control, non-target MC57 parental tumour cells ($2x10^4$ cells) as a negative control, or target MC57-HER2 tumour cells ($2x10^4$ cells) in triplicate. After 24 hours, supernatants of co-cultures were collected and used in a LEGENDplex Mouse T Helper Cytokine Panel Version 2 Flexi Kit (Biolegend) for IFN-γ, IL-2 and TNFα, and LEGENDplex Mouse Cytokine Panel 2 Flexi Kit (Biolegend) for GM-CSF according to manufacturer's instructions. All samples were analysed using a LSR II Fortessa or FACSVerse (BD Bioscience) and concentration determined against a standard curve of each analyte using FlowJo™ v10 software.

**Confocal Microscopy and Cluster analysis**

$8x10^5$ cells were labelled with unconjugated anti-Myc primary antibody (Cell signalling) in PBS/0.5% BSA for 30 minutes on ice. Cells were washed twice in PBS and further incubated with Alexafluor488 anti-mouse IgG secondary antibody (Abcam) in 50μl ice-cold RPMI for 10 minutes on ice. 50μl pre-warmed RPMI was added and samples were transferred to a 37 °C water bath for 10 min to induce CAR clustering. CAR clustering was halted via addition of ice-cold PBS/0.1 % sodium azide. Cells were washed in PBS/0.1 % sodium azide then stained on ice for 45 minutes with anti-CD28 APC (Biolegend) diluted in PBS/0.1 % sodium azide. Cells were fixed with 3% paraformaldehyde, transferred to 8-well chamber slides (Ibidi) and stored at 4°C overnight until imaging. 3D confocal image data was collected using a Zeiss LSM 980 microscope, with 55-60 slices collected per image at a z-step size of 0.23um. The pinhole size used was 1 airy unit resulting in a slice thickness of 600nm.

Image analysis was conducted using the cluster-picking function within the Imaris software package. CAR clusters (Alexafluor488) and CD28 clusters (APC) were counted, with the percentage of CAR clusters co-localising with a CD28 cluster reported per cell with at least 30 cells per construct analysed.

**In vivo tumour growth**

For in vivo tumour growth analysis, $5x10^5$ MC38 colon adenocarcinoma cells stably expressing human HER2 (MC38-HER2) were injected subcutaneously into the left flank of 5-6 week old NSG mice, with 5-6 mice/group. One day later mice were injected intravenously via the tail vein with $1x10^7$ $CD8^+$ CAR-T cells. On days 1, 2 and 3, mice were injected intraperitoneally with $5x10^4$ IU recombinant human IL-2. Mice were weighed weekly and tumours measured daily until each individual tumour reached a maximum tumour volume of 1000mm$^3$ as per ethical guidelines, after which mice were euthanized.

#### RosettaScripts and commandlines

#### Fold, Dock, and Design

```
<ROSETTASCRIPTS>
  <SCOREFXNS>
    <ScoreFunction name="score0" weights="%%score_func_0%%" symmetric="1">
      <Reweight scoretype="mp_helicality" weight="100"/>
    </ScoreFunction>
    <ScoreFunction name="score1" weights="%%score_func_1%%" symmetric="1">
      <Reweight scoretype="mp_helicality" weight="100"/>
    </ScoreFunction>
    <ScoreFunction name="score2" weights="%%score_func_2%%" symmetric="1">
      <Reweight scoretype="mp_helicality" weight="100"/>
    </ScoreFunction>
    <ScoreFunction name="score3" weights="%%score_func_3%%" symmetric="1">
      <Reweight scoretype="mp_helicality" weight="100"/>
    </ScoreFunction>
    <ScoreFunction name="score5" weights="%%score_func_5%%" symmetric="1">
      <Reweight scoretype="mp_helicality" weight="100"/>
    </ScoreFunction>

    <ScoreFunction name="beta" weights="ref2015_memb" symmetric="1">
      <Reweight scoretype="mp_helicality" weight="100"/>
    </ScoreFunction>
    <ScoreFunction name="betaNotSymm" weights="ref2015_memb" symmetric="0">
      <Reweight scoretype="mp_helicality" weight="100"/>
    </ScoreFunction>

    <ScoreFunction name="helicality" symmetric="1">
      <Reweight scoretype="mp_helicality" weight="1"/>
    </ScoreFunction>
  </SCOREFXNS>
  <TASKOPERATIONS>
    <InitializeFromCommandline name="init"/>
    <RestrictToRepacking name="rtr"/>
  </TASKOPERATIONS>
  <MOVERS>
      <PyMOLMover name="pmm" keep_history="1"/>
    <SetupForSymmetry name="symm" definition="%%symm_file%%"/>
    <SymmetricAddMembraneMover name="add_memb"
membrane_core="%%membrane_core%%" steepness="%%steepness%%">
      <Span start="%%span_start_1%%" end="%%span_end_1%%"
orientation="%%span_orientation_1%%"/>
      <Span start="%%span_start_2%%" end="%%span_end_2%%"
orientation="%%span_orientation_2%%"/>
    </SymmetricAddMembraneMover>
    <MembranePositionFromTopologyMover name="init_pos"/>
    <FastRelax name="fast_relax" scorefxn="beta"/>
```

```
    Fragment movers
    <SingleFragmentMover name="frag9" fragments="%%frags9mers%%"
policy="uniform"/>
    <SingleFragmentMover name="frag3" fragments="%%frags3mers%%"
policy="smooth"/>
    Fold-and-dock specific movers
    <SymFoldandDockRbTrialMover name="rbtrial" rot_mag="8.0" trans_mag="3.0"
rotate_anchor_to_x="1"/>
    <SymFoldandDockRbTrialMover name="rbtrial_smooth" rot_mag="1.0"
trans_mag="0.1" rotate_anchor_to_x="1"/>
    <SymFoldandDockMoveRbJumpMover name="rbjump"/>
    <SymFoldandDockSlideTrialMover name="slidetrial"/>

    Random movers
    <RandomMover name="early_stage_moveset"
movers="frag9,rbtrial,rbjump,slidetrial" weights="1.0,0.2,1.0,0.1"
repeats="1"/>
    <RandomMover name="final_stage_moveset"
movers="frag3,rbtrial_smooth,rbjump,slidetrial" weights="1.0,0.2,1.0,0.1"
repeats="1"/>

    Monte Carlo Movers
    <GenericMonteCarlo name="stage1" scorefxn_name="score0"
mover_name="early_stage_moveset" temperature="2.0" trials="200"
recover_low="1"/>
    <GenericMonteCarlo name="stage2" scorefxn_name="score1"
mover_name="early_stage_moveset" temperature="2.0" trials="200"
recover_low="1"/>
    <GenericMonteCarlo name="stage3a" scorefxn_name="score2"
mover_name="early_stage_moveset" temperature="2.0" trials="20"
recover_low="1"/>
    <GenericMonteCarlo name="stage3b" scorefxn_name="score5"
mover_name="early_stage_moveset" temperature="2.0" trials="20"
recover_low="1"/>
    <GenericMonteCarlo name="stage4" scorefxn_name="score3"
mover_name="final_stage_moveset" temperature="2.0" trials="400"
recover_low="1"/>

    Special stage 3 logic
    <ParsedProtocol name="stage3_cyc">
      <Add mover="stage3a"/>
      <Add mover="stage3b"/>
    </ParsedProtocol>
    <LoopOver name="stage3" mover_name="stage3_cyc" iterations="5"
drift="1"/>

    Converts the centroid-level pose to fullatom for scoring
    <SwitchResidueTypeSetMover name="fullatom" set="fa_standard"/>
    <ExtractAsymmetricPose name="extract_asp"/>

    <!--<SymPackRotamersMover name="soft_design" scorefxn="beta_soft"
task_operations="init"/>-->
    <TaskAwareSymMinMover name="hard_min" scorefxn="beta" chi="1" bb="1"
rb="1" task_operations="init"/>
    <SymPackRotamersMover name="hard_design" scorefxn="beta"
task_operations="init"/>
```

```
    <RotamerTrialsMinMover name="RTmin" scorefxn="beta"
task_operations="init,rtr"/>
  </MOVERS>
  <FILTERS>
    <ScoreType name="total" scorefxn="beta" score_type="total_score"
confidence="0" threshold="0"/>

    <Sasa name="a_sasa" confidence="0"/>

    <ResidueLipophilicity name="a_res_solv" threshold="1000"
confidence="0"/>
    <SpanTopologyMatchPose name="a_span_topo" confidence="0"/>
    <Ddg name="a_ddg" scorefxn="betaNotSymm" chain_num="2" repeats="5"
extreme_value_removal="true" confidence="0"/>
    <PackStat name="a_pack" confidence="0"/>
    <BuriedUnsatHbonds2 name="a_unsat" scorefxn="beta" confidence="0"/>
        <ShapeComplementarity name="a_shape" confidence="0"/>
    <TMsSpanMembrane name="a_tms_span" confidence="1"/>
    <TMsSpanMembrane name="a_tms_span_fa" confidence="1" min_distance="25"/>
    <HelixHelixAngle name="a_hha_ang" angle_or_dist="angle"
start_helix_1="%%span_start_1%%" end_helix_1="%%span_end_1%%"
start_helix_2="%%span_start_2%%" end_helix_2="%%span_end_2%%"
confidence="0"/>
    <HelixHelixAngle name="a_hha_dst_vec" angle_or_dist="dist"
dist_by_atom="0" start_helix_1="%%span_start_1%%"
end_helix_1="%%span_end_1%%" start_helix_2="%%span_start_2%%"
end_helix_2="%%span_end_2%%" confidence="0"/>
    <HelixHelixAngle name="a_hha_dst_atm" angle_or_dist="dist"
dist_by_atom="1" start_helix_1="%%span_start_1%%"
end_helix_1="%%span_end_1%%" start_helix_2="%%span_start_2%%"
end_helix_2="%%span_end_2%%" confidence="0"/>
    <ScoreType name="a_helicality" scorefxn="helicality"
score_type="total_score" confidence="1" threshold="3"/>
    <TMsAAComp name="a_tms_aa_comp" confidence="0" threshold="0"/>
  </FILTERS>
  <PROTOCOLS>
   <Add mover="symm"/>
   <Add mover="add_memb"/>
   <Add mover="stage1"/>
   <Add mover="stage2"/>
   <Add mover="stage3"/>
   <Add mover="stage4"/>
   <Add filter="a_helicality"/>
   <Add mover="fullatom"/>
   <Add filter="a_tms_span_fa"/>
   !!!! design !!!!
   <Add mover="hard_design"/>
   <Add mover="hard_min"/>
   <Add mover="hard_design"/>
   <Add mover="hard_min"/>
   <Add mover="RTmin"/>
   <Add mover="RTmin"/>
   <Add filter="a_tms_span"/>
   <Add mover="fast_relax"/>
   <Add mover="pmm"/>  # DO I WANT THIS HERE?
   <Add filter="total"/>
   <Add filter="a_sasa"/>
```

```
    <Add filter="a_res_solv"/>
    <Add filter="a_span_topo"/>
    <Add filter="a_helicality"/>
    # important to remove Symmetry so RMSD can be calculated
    <Add mover="extract_asp"/>
    <Add filter="a_res_solv"/>
    <Add filter="a_pack"/>
    <Add filter="a_unsat"/>
    <Add filter="a_shape"/>
    <Add filter="a_ddg"/>
    <Add filter="a_hha_ang"/>
    <Add filter="a_hha_dst_vec"/>
    <Add filter="a_hha_dst_atm"/>
    <Add filter="a_tms_aa_comp"/>
  </PROTOCOLS>
  <OUTPUT scorefxn="betaNotSymm"/>
</ROSETTASCRIPTS>
```

**Command line**

```
~/Rosetta/main/source/bin/rosetta_scripts.default.linuxgccreleas  -database
Rosetta/main/database @flags_fdd
flags_fdd:
# general data
-parser:protocol fold_dock_design.xml
-in:file:fasta ploy_V.fasta
-in:file:native 24.pdb
-overwrite
-use_input_sc
-nstruct 12
-jd2:ntrials 10
#-mute all
# fragment stuff
-parser:script_vars frags9mers=frags_9ploy_V_pdbtm.200.9mers
-parser:script_vars frags3mers=frags_3ploy_V_pdbtm.200.3mers
-parser:script_vars symm_file=C2.symm
# membrane spans:
-parser:script_vars span_starts=1A
-parser:script_vars span_ends=24A
-parser:script_vars span_oris=out2in
-parser:script_vars span_start_1=1
-parser:script_vars span_end_1=24
-parser:script_vars span_orientation_1=out2in
-parser:script_vars span_start_2=25
-parser:script_vars span_end_2=48
-parser:script_vars span_orientation_2=out2in
-parser:script_vars span_starts=1A
-parser:script_vars span_ends=24A
-parser:script_vars span_oris=out2in
-parser:script_vars score_func_0=score0_memb
-parser:script_vars score_func_1=score1_memb
-parser:script_vars score_func_2=score2_memb
-parser:script_vars score_func_3=score3_memb
-parser:script_vars score_func_5=score5_memb
# energy function stuff:
-parser:script_vars energy_function=ref
# membrane adjustments
-mp:scoring:hbond
```

```
-parser:script_vars steepness=4
-parser:script_vars membrane_core=10
```

**Sequence diversification**

```
<ROSETTASCRIPTS>
  <SCOREFXNS>
    <ScoreFunction name="ref" weights="ref2015_memb" symmetric="1">
      <Reweight scoretype="fa_mpenv_smooth" weight="0.000001"/>
      <Reweight scoretype="coordinate_constraint" weight="0.4"/>
    </ScoreFunction>
    <ScoreFunction name="ref_no_cst" weights="ref2015_memb" symmetric="1">
      <Reweight scoretype="fa_mpenv_smooth" weight="0.000001"/>
    </ScoreFunction>
    <ScoreFunction name="helicality" symmetric="1">
      <Reweight scoretype="mp_helicality" weight="1"/>
    </ScoreFunction>
  </SCOREFXNS>
  <RESIDUE_SELECTORS>
    <ResidueName name="gly" residue_name3="GLY"/>
    <ResidueName name="asn_gln" residue_name3="ASN,GLN"/>
    <Index name="all_aas" resnums="1-48"/>
    <Not name="not_aas" selector="all_aas"/>
  </RESIDUE_SELECTORS>
  <TASKOPERATIONS>
    <OperateOnResidueSubset name="keep_gly" selector="gly">
      <RestrictToRepackingRLT/>
    </OperateOnResidueSubset>
    <OperateOnResidueSubset name="keep_qn" selector="asn_gln">
      <RestrictToRepackingRLT/>
    </OperateOnResidueSubset>
    <OperateOnResidueSubset name="freeze_not_aas" selector="not_aas">
      <PreventRepackingRLT/>
    </OperateOnResidueSubset>
    <RestrictToRepacking name="rtr"/>
    <RestrictAbsentCanonicalAAS name="to_restrict_avlimfwyst" resnum="0"
keep_aas="GAVIFSTYW"/>
    <InitializeFromCommandline name="init"/>
  </TASKOPERATIONS>
  <FILTERS>
    <ResidueLipophilicity name="a_res_lipo" confidence="0"/>
    <ScoreType name="a_total" scorefxn="ref" score_type="total_score"
confidence="0" threshold="0"/>
    <ScoreType name="a_total_final" scorefxn="ref" score_type="total_score"
confidence="1" threshold="-140"/>
    <Sasa name="a_sasa" confidence="0"/>
    <SpanTopologyMatchPose name="a_span_topo" confidence="0"/>
    <Ddg name="a_ddg" scorefxn="ref_no_cst" confidence="0" />
    <PackStat name="a_pack" confidence="1" threshold="0.3"/>
    <BuriedUnsatHbonds2 name="a_unsat" scorefxn="ref" confidence="0"/>
    <ShapeComplementarity name="a_shape" confidence="0"/>
    <TMsSpanMembrane name="a_tms_span" confidence="0"/>
    <TMsSpanMembrane name="a_tms_span_fa" confidence="0" min_distance="25"/>
    <HelixHelixAngle name="a_hha_ang" angle_or_dist="angle"
start_helix_1="%%s1%%" end_helix_1="%%e1%%" start_helix_2="%%s2%%"
end_helix_2="%%e2%%" confidence="0"/>
    <MembAccesResidueLipophilicity name="a_marl" confidence="0"
verbose="0"/>
    <TMsAAComp name="a_tms_aa_comp" confidence="0" threshold="0"/>
```

```
    <BindingStrain name="a_bind" scorefxn="ref_no_cst" jump="1"
confidence="1" threshold="5"/>
    <ScoreType name="a_helicality" scorefxn="helicality"
score_type="total_score" confidence="1" threshold="3"/>
    <Sigmoid name="a_total_sig" filter="a_total" steepness="0.5" offset="20"
negate="0"  confidence="0"/>
    <Sigmoid name="a_ddg_sig_10" filter="a_ddg" steepness="1" offset="10"
negate="0" confidence="0"/>
    <Sigmoid name="a_ddg_sig_2" filter="a_ddg" steepness="1" offset="3"
negate="0" confidence="0"/>
    <Sigmoid name="a_comp_sig" filter="a_tms_aa_comp" steepness="50"
offset="0.05" negate="0" confidence="0"/>
    <Operator name="a_obj_func_ddg_10"
filters="a_total_sig,a_ddg_sig_10,a_comp_sig" operation="PRODUCT"
logarithm="1" threshold="100000" confidence="0" negate="1"/>
    <Operator name="a_obj_func_ddg_2"
filters="a_total_sig,a_ddg_sig_2,a_comp_sig" operation="PRODUCT"
logarithm="1" threshold="100000" confidence="0" negate="1"/>
  </FILTERS>
  <MOVERS>
    <SymmetricAddMembraneMover name="add_memb" membrane_core="%%memb_core%%"
steepness="%%steepness%%">
      <Span start="%%s1%%" end="%%e1%%" orientation="%%o1%%"/>
      <Span start="%%s2%%" end="%%e2%%" orientation="%%o2%%"/>
    </SymmetricAddMembraneMover>
    <RandomMutation name="mutate"
task_operations="freeze_not_aas,init,to_restrict_avlimfwyst,keep_gly,keep_qn
" scorefxn="ref"/>
    <VirtualRoot name="virt_root"/>
    <AtomCoordinateCstMover name="atom_cst" coord_dev="0.5" bounded="false"
native="false"/>
    <TaskAwareSymMinMover name="min_mover" scorefxn="ref" chi="1" bb="1"
rb="1" task_operations="init,rtr"/>
    <ParsedProtocol name="mutate_min">
      <Add mover="mutate"/>
      <Add mover="min_mover"/>
    </ParsedProtocol>
    <GenericMonteCarlo name="gmc_sigs_1" filter_name="a_obj_func_ddg_10"
preapply="0" mover_name="mutate_min" temperature="0.1" trials="240"
recover_low="1" reset_baselines="1"/>
    <GenericMonteCarlo name="gmc_sigs_2" filter_name="a_obj_func_ddg_2"
preapply="0" mover_name="mutate_min" temperature="0.1" trials="80"
recover_low="1" reset_baselines="1"/>
    <GenericSimulatedAnnealer name="sim_anneal" mover_name="mutate_min"
filter_name="a_obj_func_ddg_2" trials="120" sample_type="low"
recover_low="1" preapply="0" reset_baselines="1" history="10"/>
    <SetupForSymmetry name="symm" definition="%%symm_file%%"
preserve_datacache="false"/>
  </MOVERS>
  <PROTOCOLS>
    <Add mover="symm"/>
    <Add mover="add_memb"/>
    <Add mover="atom_cst"/>
    <Add mover="min_mover"/>
    <Add mover="sim_anneal"/>
    <Add mover="min_mover"/>
    <Add filter="a_total_sig"/>
```

```
    <Add filter="a_total_final"/>
    <Add filter="a_ddg_sig_2"/>
    <Add filter="a_ddg_sig_10"/>
    <Add filter="a_comp_sig"/>
    <Add filter="a_obj_func_ddg_2"/>
    <Add filter="a_obj_func_ddg_10"/>
    <Add filter="a_tms_aa_comp"/>
    <Add filter="a_res_lipo"/>
    <Add filter="a_marl"/>
    <Add filter="a_sasa"/>
    <Add filter="a_total"/>
    <Add filter="a_ddg"/>
    <Add filter="a_bind"/>
    <Add filter="a_pack"/>
    <Add filter="a_unsat"/>
    <Add filter="a_shape"/>
    <Add filter="a_tms_span"/>
    <Add filter="a_helicality"/>
    <Add filter="a_tms_aa_comp"/>
  </PROTOCOLS>
  <OUTPUT scorefxn="ref"/>
</ROSETTASCRIPTS>
```

**Command line**

```
~/Rosetta/main/source/bin/rosetta_scripts.default.linuxgccreleas -database
Rosetta/main/database @flags_seq_divers
```

```
flags_seq_divers:
-parser:protocol GMC_seq_diversifier.xml
-overwrite
-parser:script_vars memb_core=10
-parser:script_vars steepness=4
-mute all
-nstruct 10
-parser:script_vars s1=1
-parser:script_vars s2=24
-parser:script_vars e1=25
-parser:script_vars e2=48
-parser:script_vars o1=out2in
-parser:script_vars o2=out2in
-mp:scoring:hbond
-jd2:ntrials 1000000
-use_input_sc
-s seed_INPUT.pdb
-parser:script_vars symm_file=seed.symm
```

**FilterScan**

```
<ROSETTASCRIPTS>
    <SCOREFXNS>
        <ScoreFunction name="ddg_sfx" weights="%%scorefxn%%" symmetric="1"/>
        <ScoreFunction name="full" weights="ref2015_memb" symmetric="1">
          <Reweight scoretype="coordinate_constraint"
weight="%%cst_value%%"/>
        </ScoreFunction>
        <ScoreFunction name="soft" weights="ref2015_soft" symmetric="1">
          <Reweight scoretype="mp_res_lipo" weight="1"/>
          <Reweight scoretype="coordinate_constraint"
weight="%%cst_value%%"/>
        </ScoreFunction>
    </SCOREFXNS>
```

```
    <TASKOPERATIONS>
        <InitializeFromCommandline name="init"/>
       <RestrictToRepacking name="rtr"/>
      <DesignAround name="des_around" design_shell="0.1"
resnums="%%current_res%%" repack_shell="8.0"/>
        <OperateOnResidueSubset name="restrict_res">
            <Index resnums="%%res_to_restrict%%"/>
            <RestrictToRepackingRLT/>
        </OperateOnResidueSubset>
        <OperateOnResidueSubset name="fix_res">
            <Index resnums="%%res_to_fix%%"/>
            <PreventRepackingRLT/>
        </OperateOnResidueSubset>
    </TASKOPERATIONS>
    <MOVERS>
      <SetupForSymmetry name="symm" definition="%%symm_file%%"/>
      <SymmetricAddMembraneMover name="add_memb" membrane_core="10"
steepness="4">
        <Span start="1" end="24" orientation="in2out"/>
        <Span start="25" end="48" orientation="in2out"/>
</SymmetricAddMembraneMover>
      <TransformIntoMembraneMover name="tramsform" />
      <AtomCoordinateCstMover name="atom_cst" coord_dev="0.5"
bounded="false" native="false"/>
      <MinMover name="min_all" scorefxn="ddg_sfx" chi="1" bb="1"
jump="%%jump%%"/>#scorefxn_full
    <SymPackRotamersMover name="soft_repack" scorefxn="soft"
task_operations="init,rtr"/>
    <SymPackRotamersMover name="hard_repack" scorefxn="full"
task_operations="init,rtr"/>
    <SymRotamerTrialsMover name="RTmin" scorefxn="full"
task_operations="init,rtr"/>
    <SymMinMover name="soft_min" scorefxn="soft" chi="1" bb="1" jump="0"/>
    <SymMinMover name="hard_min" scorefxn="full" chi="1" bb="1" jump="0"/>
    <ParsedProtocol name="refinement_block"> #10 movers
      <Add mover_name="soft_repack"/>
      <Add mover_name="soft_min"/>
      <Add mover_name="soft_repack"/>
      <Add mover_name="hard_min"/>
      <Add mover_name="hard_repack"/>
      <Add mover_name="hard_min"/>
      <Add mover_name="hard_repack"/>
      Add mover_name="RTmin"/>
      Add mover_name="RTmin"/>
      <Add mover_name="hard_min"/>
    </ParsedProtocol>
    <LoopOver name="iter4" mover_name="refinement_block" iterations="4"/>

    </MOVERS>

    <FILTERS>
         <Ddg name="ddg" scorefxn="ddg_sfx" threshold="0"
repeats="5"/>#chain_num="2"
        <ScoreType name="stability_score_full" scorefxn="full"
score_type="total_score" threshold="0.0"/>
```

```
      <Delta name="delta_score_full" filter="stability_score_full" upper="1"
lower="0" range="0.5"/> #upper and lower are booleans. Delta filters out all
the mutations that are worse or better by less than -0.55R.E.U
        Delta name="delta_score_full" filter="stability_score_full"
upper="1" lower="0" range="0.5"/> #upper and lower are booleans. Delta
filters out all the mutations that are worse or better by less than
-0.55R.E.U

              FilterScan name="filter_scan" scorefxn="ddg_sfx"
relax_mover="min_all" keep_native="%%keep_n%%"
task_operations="init,des_around,fix_res,restrict_res"
delta_filters="delta_score_full" delta="true"
resfile_name="%%resfiles_path%%/res_%%current_res%%" report_all="1"
delta_filter_thresholds="%%fs_thresholds%%"
score_log_file="%%scores_path%%/res%%current_res%%_score_full.log"
dump_pdb="1" />
             <FilterScan name="filter_scan" scorefxn="full"
relax_mover="iter4" keep_native="%%keep_n%%"
task_operations="init,des_around,fix_res,restrict_res"
delta_filters="delta_score_full" delta="true"
resfile_name="%%resfiles_path%%/res_%%current_res%%" report_all="1"
delta_filter_thresholds="%%fs_thresholds%%"
score_log_file="%%scores_path%%/res%%current_res%%_score_full.log"
dump_pdb="0" />
    </FILTERS>
    <PROTOCOLS>
              <Add mover="symm"/>
           <Add mover="add_memb"/>
           Add mover="tramsform"/>
        <Add mover="atom_cst"/>
        <Add filter="filter_scan"/>
    </PROTOCOLS>
</ROSETTASCRIPTS>
```
**Command line**
```
~/Rosetta/main/source/bin/rosetta_scripts.default.linuxgccreleas -database
Rosetta/main/database @flags_filterscan
Flags_filterscan:
# general data
-parser:protocol filterscan_auto_refine_SYMM.xml
#-database #path to database
-overwrite
# membrane spans: #changed to in2out 23Feb17 for experimental reasons
-parser:script_vars res_to_fix=1A
-parser:script_vars res_to_restrict=1A
-parser:script_vars cst_value=0.4
-parser:script_vars jump=1
-parser:script_vars scorefxn=ref2015_memb
-parser:script_vars span_orientation_2=in2out
-parser:script_vars
fs_thresholds=0.0,0.5,1.0,1.5,2.0,2.5,3.0,3.5,4.0,4.5,5.0,5.5,6.0,6.5,7.0,8.
0,9.0,10.0,11.0,12.0
-parser:script_vars keep_n=1
#-score::elec_memb_sig_die
#-corrections::beta_nov16
#-score:memb_fa_sol
-mp:scoring:hbond
-use_input_sc
```

```
-parser:script_vars  current_res=3
-parser:script_vars pdb_dump=#filterscan/pdbs/3_
-out:path:score  score
-out:path:pdb  pdbs
-parser:script_vars  resfiles_path=./
-parser:script_vars  scores_path=./
-parser:script_vars  symm_file=seed.symm
```

#### ***ab-initio*** **structure prediction**

```
<ROSETTASCRIPTS>
  <TASKOPERATIONS>
    <InitializeFromCommandline name="init"/>
    <RestrictToRepacking name="rtr"/>
  </TASKOPERATIONS>
  <SCOREFXNS>
    <ScoreFunction name="score0" weights="%%score_func_0%%" symmetric="1">
      <Reweight scoretype="mp_helicality" weight="0"/>
        <Reweight scoretype="mp_span_ang" weight="0"/>
    </ScoreFunction>
    <ScoreFunction name="score1" weights="%%score_func_1%%" symmetric="1">
      <Reweight scoretype="mp_helicality" weight="0"/>
        <Reweight scoretype="mp_span_ang" weight="0"/>
        <Reweight scoretype="mp_nonhelix" weight="0"/>
    </ScoreFunction>
    <ScoreFunction name="score2" weights="%%score_func_2%%" symmetric="1">
      <Reweight scoretype="mp_helicality" weight="0"/>
        <Reweight scoretype="mp_span_ang" weight="0"/>
        <Reweight scoretype="mp_nonhelix" weight="0"/>
    </ScoreFunction>
    <ScoreFunction name="score3" weights="%%score_func_3%%" symmetric="1">
      <Reweight scoretype="mp_helicality" weight="0"/>
        <Reweight scoretype="mp_span_ang" weight="0"/>
        <Reweight scoretype="mp_nonhelix" weight="0"/>
    </ScoreFunction>
    <ScoreFunction name="score5" weights="%%score_func_5%%" symmetric="1">
      <Reweight scoretype="mp_helicality" weight="0"/>
        <Reweight scoretype="mp_span_ang" weight="0"/>
        <Reweight scoretype="mp_nonhelix" weight="0"/>
    </ScoreFunction>

    <ScoreFunction name="ref" weights="ref2015_memb" symmetric="1">
      <Reweight scoretype="mp_helicality" weight="0"/>
        <Reweight scoretype="mp_span_ang" weight="0"/>
    </ScoreFunction>
    <ScoreFunction name="refNotSymm" weights="ref2015_memb" symmetric="0">
      <Reweight scoretype="mp_helicality" weight="0"/>
        <Reweight scoretype="mp_span_ang" weight="0"/>
    </ScoreFunction>

    <ScoreFunction name="helicality" symmetric="1">
      <Reweight scoretype="mp_helicality" weight="0"/>
        <Reweight scoretype="mp_span_ang" weight="0"/>
    </ScoreFunction>
    <ScoreFunction name="helicality_notsymm" symmetric="0">
      <Reweight scoretype="mp_helicality" weight="0"/>
        <Reweight scoretype="mp_span_ang" weight="0"/>
```

```
    </ScoreFunction>
  </SCOREFXNS>
  <MOVERS>
    <SetupForSymmetry name="symm" definition="%%symm_file%%"/>
    <SymmetricAddMembraneMover name="add_memb"
membrane_core="%%membrane_core%%" steepness="%%steepness%%"
span_starts_num="%%span_starts%%" span_ends_num="%%span_ends%%"
span_orientations="%%span_oris%%"/>
    <MembranePositionFromTopologyMover name="init_pos"/>
    <FastRelax name="fast_relax" scorefxn="%%energy_function%%"
task_operations="init"/>

    Fragment movers
    <SingleFragmentMover name="frag9" fragments="%%frags9mers%%"
policy="uniform">
      <MoveMap>
        <Span begin="1" end="24" chi="1" bb="1"/>
      </MoveMap>
    </SingleFragmentMover>
    <SingleFragmentMover name="frag3" fragments="%%frags3mers%%"
policy="smooth">
      <MoveMap>
        <Span begin="1" end="24" chi="1" bb="1"/>
      </MoveMap>
    </SingleFragmentMover>

    Fold-and-dock specific movers
    <SymFoldandDockRbTrialMover name="rbtrial" rot_mag="8.0" trans_mag="3.0"
rotate_anchor_to_x="1"/>
    <SymFoldandDockRbTrialMover name="rbtrial_smooth" rot_mag="1.0"
trans_mag="0.1" rotate_anchor_to_x="1"/>
    <SymFoldandDockMoveRbJumpMover name="rbjump"/>
    <SymFoldandDockSlideTrialMover name="slidetrial"/>

    Random movers
    <RandomMover name="early_stage_moveset"
movers="frag9,rbtrial,rbjump,slidetrial" weights="1.0,0.2,1.0,0.1"
repeats="1"/>
    <RandomMover name="final_stage_moveset"
movers="frag3,rbtrial_smooth,rbjump,slidetrial" weights="1.0,0.2,1.0,0.1"
repeats="1"/>

    Monte Carlo Movers
    <GenericMonteCarlo name="stage1" scorefxn_name="score0"
mover_name="early_stage_moveset" temperature="2.0" trials="200"
recover_low="1"/>
    <GenericMonteCarlo name="stage2" scorefxn_name="score1"
mover_name="early_stage_moveset" temperature="2.0" trials="200"
recover_low="1"/>
    <GenericMonteCarlo name="stage3a" scorefxn_name="score2"
mover_name="early_stage_moveset" temperature="2.0" trials="20"
recover_low="1"/>
    <GenericMonteCarlo name="stage3b" scorefxn_name="score5"
mover_name="early_stage_moveset" temperature="2.0" trials="20"
recover_low="1"/>
```

```
    <GenericMonteCarlo name="stage4" scorefxn_name="score3"
mover_name="final_stage_moveset" temperature="2.0" trials="400"
recover_low="1"/>

    Special stage 3 logic
    <ParsedProtocol name="stage3_cyc">
      <Add mover="stage3a"/>
      <Add mover="stage3b"/>
    </ParsedProtocol>
    <LoopOver name="stage3" mover_name="stage3_cyc" iterations="5"
drift="1"/>

    Converts the centroid-level pose to fullatom for scoring
    <SwitchResidueTypeSetMover name="fullatom" set="fa_standard"/>
    <ExtractAsymmetricPose name="extract_asp" clear_sym_def="1"/>
    <MinMover name="min_mover" scorefxn="refNotSymm" chi="1" bb="1"
jump="1"/>
    <PackRotamersMover name="pack" scorefxn="refNotSymm"
task_operations="init,rtr"/>
    <RotamerTrialsMinMover name="RTmin" scorefxn="refNotSymm"
task_operations="init,rtr"/>
    <DumpPdb name="dump_pdb" fname="dump.pdb"
scorefxn="%%energy_function%%"/>

    <SwitchChainOrder name="switch" chain_order="123"/>
    <DeleteChain name="delete_mem" chain="4" />

  </MOVERS>
  <FILTERS>
    <ScoreType name="total" scorefxn="%%energy_function%%"
score_type="total_score" confidence="1" threshold="0"/>
    <Sasa name="a_sasa" confidence="0" threshold="300"/>

    <ResidueLipophilicity name="a_res_lipo" threshold="1000"
confidence="0"/>
    <SpanTopologyMatchPose name="a_span_topo" confidence="0"/>
    <Ddg name="a_ddg" scorefxn="%%energy_function%%NotSymm" chain_num="2"
repeats="5" extreme_value_removal="true" confidence="0" threshold="-5"/>
    <PackStat name="a_pack" confidence="0" threshold="0.3"/>
    <BuriedUnsatHbonds2 name="a_unsat" scorefxn="%%energy_function%%"
confidence="0"/>
    <ShapeComplementarity name="a_shape" confidence="0"/>
    <TMsSpanMembrane name="a_tms_span" confidence="1" min_distance="25"/>
    <TMsSpanMembrane name="a_tms_span_fa" confidence="0" min_distance="25"/>
    <HelixHelixAngle name="a_hha_ang" angle_or_dist="angle"
start_helix_1="%%span_start_1%%" end_helix_1="%%span_end_1%%"
start_helix_2="%%span_start_2%%" end_helix_2="%%span_end_2%%"
confidence="0"/>
    <HelixHelixAngle name="a_hha_dst_vec" angle_or_dist="dist"
dist_by_atom="0" start_helix_1="%%span_start_1%%"
end_helix_1="%%span_end_1%%" start_helix_2="%%span_start_2%%"
end_helix_2="%%span_end_2%%" confidence="0"/>
    <HelixHelixAngle name="a_hha_dst_atm" angle_or_dist="dist"
dist_by_atom="1" start_helix_1="%%span_start_1%%"
end_helix_1="%%span_end_1%%" start_helix_2="%%span_start_2%%"
end_helix_2="%%span_end_2%%" confidence="0"/>
```

```
    <MembAccesResidueLipophilicity name="a_marl" confidence="0" 
verbose="0"/>
    <ScoreType name="a_helicality" scorefxn="helicality_notsymm" 
score_type="mp_helicality" confidence="0" threshold="10"/>
    <ScoreType name="a_helicality_symm" scorefxn="helicality" 
score_type="mp_helicality" confidence="0" threshold="10"/>
    <MPSpanAngle name="a_angle_1" tm="1" ang_min="0" ang_max="50" 
confidence="0"/>
    <MPSpanAngle name="a_angle_2" tm="2" ang_min="0" ang_max="50" 
confidence="0"/>

    <BindingStrain name="a_bind" scorefxn="%%energy_function%%" jump="1" 
confidence="0" threshold="5"/>
    <PoseInfo name="info"/>
  </FILTERS>
  <PROTOCOLS>
   <Add mover="symm"/>
   <Add mover="add_memb"/>

   <Add mover="stage1"/>
   <Add mover="stage2"/>
   <Add mover="stage3"/>
   <Add mover="stage4"/>

   <Add filter="a_helicality_symm"/>
   <Add filter="a_angle_1"/>
   <Add filter="a_angle_2"/>

   <Add mover="fullatom"/>

   <Add filter="a_tms_span"/>
   <Add mover="fast_relax"/>

   <Add filter="total"/>
   <Add filter="a_sasa"/>

   <Add filter="a_span_topo"/>

   <Add mover="extract_asp"/>
   <Add mover="pack"/>
   <Add mover="min_mover"/>
   <Add mover="RTmin"/>
   <Add mover="RTmin"/>

   <Add filter="a_tms_span"/>
   <Add filter="total"/>
   <Add filter="a_sasa"/>
   <Add filter="a_span_topo"/>

   <Add filter="a_res_lipo"/>
   <Add filter="a_pack"/>
   <Add filter="a_unsat"/>
   <Add filter="a_shape"/>
   <Add filter="a_ddg"/>
   <Add filter="a_hha_ang"/>
   <Add filter="a_hha_dst_vec"/>
   <Add filter="a_hha_dst_atm"/>
```

```
    <Add filter="a_marl"/>
    <Add filter="a_tms_span_fa"/>
    <Add filter="a_helicality"/>
    <Add filter="a_angle_1"/>
    <Add filter="a_angle_2"/>
    <Add filter="a_bind"/>
  </PROTOCOLS>
  <OUTPUT scorefxn="%%energy_function%%NotSymm"/>
</ROSETTASCRIPTS>
```

**Methods references**


Das, R., André, I., Shen, Y., Wu, Y., Lemak, A., Bansal, S., Arrowsmith, C.H., Szyperski, T., and Baker, D. (2009). Simultaneous prediction of protein folding and docking at high resolution. Proceedings of the National Academy of Sciences *106*, 18978–18983.

Duong, M.T., Jaszewski, T.M., Fleming, K.G., and MacKenzie, K.R. (2007). Changes in apparent free energy of helix-helix dimerization in a biological membrane due to point mutations. J. Mol. Biol. *371*, 422–434.

Elazar, A., Weinstein, J., Biran, I., Fridman, Y., Bibi, E., and Fleishman, S.J. (2016). Mutational scanning reveals the determinants of protein insertion and association energetics in the plasma membrane. Elife *5*, 12125.

Emsley, P., Lohkamp, B., Scott, W.G., and Cowtan, K. (2010). Features and development of Coot. Acta Crystallogr. D Biol. Crystallogr. *66*, 486–501.

Fleishman, S.J., Leaver-Fay, A., Corn, J.E., Strauch, E.M., Khare, S.D., Koga, N., Ashworth, J., Murphy, P., Richter, F., Lemmon, G., et al. (2011). Rosettascripts: A scripting language interface to the Rosetta Macromolecular modeling suite. PLoS One *6*, e20161.

Gront, D., Kulp, D.W., Vernon, R.M., Strauss, C.E.M., and Baker, D. (2011). Generalized fragment picking in rosetta: Design, protocols and applications. PLoS One *6*.

Haynes, N.M., Trapani, J.A., Teng, M.W.L., Jackson, J.T., Cerruti, L., Jane, S.M., Kershaw, M.H., Smyth, M.J., and Darcy, P.K. (2002). Single-chain antigen recognition receptors that costimulate potent rejection of established experimental tumors. Blood *100*, 3155–3163.

Kabsch, W. (2010). XDS. Acta Crystallogr. D Biol. Crystallogr. *66*, 125–132.

Langosch, D., Brosig, B., Kolmar, H., and Fritz, H.J. (1996). Dimerisation of the glycophorin A transmembrane segment in membranes probed with the ToxR transcription activator. J. Mol. Biol. *263*, 525–530.

Lawrence, M.C., and Colman, P.M. (1993). Shape complementarity at protein/protein interfaces. J. Mol. Biol. *234*, 946–950.

Liebschner, D., Afonine, P.V., Baker, M.L., Bunkóczi, G., Chen, V.B., Croll, T.I., Hintze, B., Hung, L.W., Jain, S., McCoy, A.J., et al. (2019). Macromolecular structure determination using X-rays, neutrons and electrons: recent developments in Phenix. Acta Crystallogr D Struct Biol *75*, 861–877.

Liu, Y., Engelman, D.M., and Gerstein, M. (2002). Genomic analysis of membrane protein families: abundance and conserved motifs. Genome Biol. *3*, research0054.

McCoy, A.J., Grosse-Kunstleve, R.W., Adams, P.D., Winn, M.D., Storoni, L.C., and Read, R.J. (2007). Phaser crystallographic software. J. Appl. Crystallogr. *40*, 658–674.

Russ, W.P., and Engelman, D.M. (1999). TOXCAT: a measure of transmembrane helix association in a biological membrane. Proc. Natl. Acad. Sci. U. S. A. *96*, 863–868.

Schanzenbach, C., Schmidt, F.C., Breckner, P., Teese, M.G., and Langosch, D. (2017). Identifying ionic interactions within a membrane using BLaTM, a genetic tool to measure homo- and heterotypic transmembrane helix-helix interactions. Sci. Rep. *7*, 43476.

Sharma, P., Kaywan-Lutfi, M., Krshnan, L., Byrne, E.F.X., Call, M.J., and Call, M.E. (2013). Production of disulfide-stabilized transmembrane peptide complexes for structural studies. J. Vis. Exp. e50141.

Srinivasan, S., Deng, W., and Li, R. (2011). L-selectin transmembrane and cytoplasmic domains are monomeric in membranes. Biochimica et Biophysica Acta - Biomembranes *1808*, 1709–1715.

Trenker, R., Call, M.E., and Call, M.J. (2015). Crystal Structure of the Glycophorin A Transmembrane Dimer in Lipidic Cubic Phase. J. Am. Chem. Soc. *137*, 15676–15679.

Warszawski, S., Netzer, R., Tawfik, D.S., and Fleishman, S.J. (2014). A "fuzzy"-logic language for encoding multiple physical traits in biomolecules. J. Mol. Biol. *426*, 4125–4138.

Weinstein, J.Y., Elazar, A., and Fleishman, S.J. (2019). A lipophilicity-based energy function for membrane-protein modelling and design. PLoS Comput. Biol. *15*, e1007318.

Winn, M.D., Ballard, C.C., Cowtan, K.D., Dodson, E.J., Emsley, P., Evans, P.R., Keegan, R.M., Krissinel, E.B., Leslie, A.G.W., McCoy, A., et al. (2011). Overview of the CCP4 suite and current developments. Acta Crystallogr. D Biol. Crystallogr. *67*, 235–242.
